# Pulmonary Arterial Hypertension Induces a Metabolic and Inflammatory Hepatopathy

**DOI:** 10.64898/2026.03.16.712114

**Authors:** Madelyn J. Blake, Sally E. Prins, Jeffrey C. Blake, Lynn M. Hartweck, Jenna B. Mendelson, Steeve Provencher, Sandra Breuils-Bonnet, Sébastien Bonnet, Kurt W. Prins

## Abstract

Right ventricular failure (RVF) is a robust predictor of mortality in pulmonary arterial hypertension (PAH); however, the mechanisms linking RVF to end-organ dysfunction remain unclear. Hepatic impairments portend poor outcomes in PAH, but the cell-specific effects of PAH on the human liver are unknown. Here, we performed single nucleus RNA sequencing on autopsy-derived liver tissue from five PAH patients and four non-PAH controls and compared these findings to non-alcoholic steatohepatitis (NASH) and Fontan-associated liver disease (FALD). PAH hepatocytes were characterized by a pro-proliferative, Warburg-like metabolic phenotype. PAH endothelial cells (ECs) also adopted a Warburg-like profile. Although EC PI3K-Akt activation was present in PAH and FALD ECs, only PAH ECs demonstrated impaired adhesion/barrier signaling. In PAH hepatic stellate cells (HSCs), PI3K-Akt signaling was enriched, while NASH and FALD HSCs co-activated PI3K-Akt and TGF-β. Activated HSC abundances were increased in PAH livers and associated with heightened central vein fibrosis. PAH and NASH macrophages showed elevated complement signaling but reduced JAK-STAT activity. PAH livers exhibited dysregulated vasoactive gene expression, increased interleukin-6 expression in HSCs, and suppressed hepatocyte ketone metabolism. Correlational analysis demonstrated that HSC HIF-1 activation was associated with PAH severity. In total, these findings define the metabolic and inflammatory hepatopathy of PAH.

## Introduction

Right ventricular failure (RVF) is the leading cause of death in pulmonary arterial hypertension (PAH) (1). RVF remains difficult to treat because it results in widespread end-organ compromise as demonstrated by alterations in brain, skeletal muscle, gastrointestinal tract, kidney, and liver function (2). Thus, defining and understanding how RVF impacts each of these important organs will be a key step towards improving patient outcomes.

Recent clinical data in PAH are defining the detrimental consequences of RVF on hepatic structure and function, and demonstrating liver dysfunction robustly predicts outcomes. PAH patients exhibit elevated liver fibrosis (3) and heightened liver stiffness (4), both of which correlate with RVF severity. Additionally, changes in liver function are highly prognostic in PAH as even minor perturbations in serum liver enzymes (5) (aspartate aminotransferase, alanine aminotransferase, alkaline phosphatase) or liver functional assessments (bilirubin, albumin, INR) (6) are strongly associated with adverse outcomes. Clearly, hepatic dysfunction is a robust and sensitive marker of RVF, but the cellular and molecular derangements present in PAH livers are undefined.

The prevailing theory in PAH is that hepatopathy is caused by chronic venous stasis from RVF (2); however, PAH patients frequently exhibit systemic metabolic derangements that could also contribute. At present, a comparison of PAH hepatopathy with non-alcoholic steatohepatitis (NASH), a metabolically driven liver disease, and Fontan-associated liver disease (FALD), a hepatopathy defined by chronic venous stasis, is lacking. To address this critical knowledge gap, we generated a single-cell transcriptomic atlas of PAH hepatopathy by performing single nucleus RNA sequencing (snRNAseq) of control (n=4 patients, 53,057 nuclei) and PAH (n=5 patients, 71,792 nuclei) liver samples. We then compared our findings with publicly available snRNA-seq datasets from patients with NASH and FALD to delineate phenotypic similarities and differences between PAH-associated liver disease and hepatopathies driven by systemic metabolic alterations and chronic venous stasis.

## Materials and Methods

### Sex as a Biological Variable

Regarding biological sex of the non-PAH control samples, 3 female and 1 male patient were included. PAH samples consisted of 2 males and 3 females. For the PAH samples, warm autopsies performed by IUCPQ pathology department confirmed PAH as the cause of death. Thus, our study included both male and female patients, and similar findings were reported for both sexes. Clinical metadata for these patients, including biological sex, are provided in **Supplemental Table 1**.

### Human Tissue Collection

Human control and PAH liver tissue samples were collected at Laval University (CER20773). All tissues were autopsy-derived and stored at -80C prior to analysis.

### Evaluation of Total and Perivascular Fibrosis

10-μm paraffin-embedded liver sections were stained with Masson Trichrome by the Medical University of South Carolina Histology and Immunohistochemistry Lab. The whole section image was collected on a Zeiss Airyscan Microscope and total section fibrosis was blindly quantified on ImageJ by SEP. For perivascular fibrosis, central veins from the sections were cropped and the percentage of area positive for fibrosis was blindly determined by SEP using ImageJ.

### Immunohistochemical Assessment of Smooth Muscle Actin and CD206

To evaluate smooth muscle actin immunoreactivity and CD206-positive macrophage abundance, liver sections were deparaffinized using xylene and subsequent incubations with 100% ethanol, 95% ethanol, and 70% ethanol. Slides were then placed in a water bath with 10% decloaking solution. Slides were then blocked with 5% goat serum before incubation with alpha-smooth muscle actin antibody (1A4) (Invitrogen, 53-9760-82) and CD206/MRC1(E2L9N) rabbit monoclonal antibody (Cell Signaling, 91992) overnight at 4 °C. Following primary antibody incubation, sections were blocked with 5% goat serum. Secondary antibody and 0.1% Hoechst stain incubation was performed for 30 min at 37 °C.

Sections were washed with PBS before treatment with an autofluorescence quenching kit and mounted in anti-fade reagent. Confocal micrographs were obtained on a Zeiss LSM900 Airyscan 2 confocal microscope under identical conditions. Quantification of total alpha-smooth actin area in 4-5 central veins per patient sample was blindly quantified by SEP using FIJI. The median alpha-smooth positive area was used as a single data point per patient. The number of CD206 positive cells was blindly determined by SEP from 4 discrete images per section. Each sample was assigned a median number of macrophages per section.

### Nuclei Isolation

Nuclei were extracted from 5 non-PAH control and 5 PAH patients using the Singulator S200 system (S2 Genomics, CA USA) with a modified low-volume nuclear dissociation protocol (7). Per sample, two NIC+ cartridges were used with ∼20 mg of tissue/cartridge. Following isolation, nuclei suspensions were cleaned twice using density sucrose gradient centrifugations (Sigma Nuc201) and pelleted in 1% BSA (Miltenyi 130091376). Additional purification steps were performed with propidium iodide staining and subsequent fluorescence-activated cell sorting at University of Minnesota Flow Cytometry Resource. To prevent RNA degradation, all solutions were supplemented with RNase inhibitor (0.2 U/L, Millipore Sigma 03335402001). 10x Genomics, library preparation, sequencing, and alignment to the human genome (GRCh38) were completed by the University of Minnesota Genomics Center.

### Single Nucleus RNA Sequencing of PAH, NASH, and FALD Datasets

snRNA-seq analysis was performed using RStudio v4.4. Raw data for the NASH (Wang et al.; GEO: GSE212837 (8)) and FALD (Hu et al.; GEO: GSE223843 (9)) datasets were obtained from the NIH Gene Expression Omnibus. Matrix files from CellRanger were converted into Seurat objects and subsequently processed using Seurat v5 (10). For the PAH dataset, nuclei containing between 200-8500 genes (200 < nFeature_RNA < 8500) and <5% mitochondrial DNA were included for downstream analysis. One control sample from the PAH dataset was removed following pre-processing due to low nuclei counts (n=437). For the NASH dataset, nuclei containing between 200-8000 genes (200 < nFeature_RNA < 8000) and <10% mitochondrial DNA were included for downstream analysis. For the FALD dataset, nuclei containing between 800-20000 genes (800 < nFeature_RNA < 20000) and <10% mitochondrial DNA were used for downstream analysis. Identification and removal of potential doublets within each sample was performed using DoubletFinder (11). The resulting Seurat objects were then subjected to additional processing consisting of normalization, feature selection, scaling, dimensionality reduction, and clustering, in accordance with the standard Seurat workflow (10). Azimuth was used to ensure independent clustering of different cell types without over-resolution (12). We initially identified 23, 13, and 14 unique clusters in the PAH, NASH, and FALD datasets, respectively, before determining cell types at the cluster level. Cell type annotations were assigned by identifying the highly expressed genes in each cluster via the FindConservedMarkers function in Seurat. Cellular annotations were cross-validated using the Human Protein Atlas (13), a database of established cell marker genes, for up to 20 of the most highly expressed genes in each cluster. A complete list of the cell marker genes used for cluster annotation across the three datasets is provided in **Supplemental Table 2.** Notably, two clusters in the PAH dataset and one cluster in each of the NASH and FALD datasets did not contain genes highly expressed in any cell type and thus were subsequently excluded from further analysis. A full schematic representation of the snRNA-seq workflow is provided in **Supplemental Figure 1**. In addition, violin plots quantifying RNA counts, number of genes identified, percent mitochondrial DNA, and percent ribosomal DNA for each dataset are shown in **Supplemental Figure 2**.

### Differential Gene Expression (DEG) and Pathway Enrichment Analysis

Due to observed differences in nuclei quality among the PAH, NASH, and FALD datasets, different approaches for DEG analysis were applied. For the PAH dataset, the processed nuclei were pseudobulked for DEG analysis using DESeq2 (14). Genes were considered differentially expressed if the |log2FoldChange| ≥ 0.25 and adjusted *p*-value was <0.05. For the NASH and FALD datasets, DEG analysis was performed on processed, but not pseudobulked, nuclei via the *FindMarkers* function in Seurat using a Wilcoxon rank-sum test. Genes were considered differentially expressed if |log2FoldChange| ≥ 0.25 and adjusted *p*-value was <0.05. We then performed a pathway analysis for significant DEGs from each cell type using ShinyGO (https://bioinformatics.sdstate.edu/go/), and pathways were shown if two or more pathways were enriched (15).

### Calculation of Module Scores

Module scores were computed via the *AddModuleScore* function in Seurat to assess relative pathway activity at the single-cell level. Predefined gene sets representing specific biological pathways were compiled using either the Kyoto Encyclopedia of Genes and Genomes (KEGG) or WikiPathways databases and subsequently used as input. Resulting scores were extracted from the Seurat objects and then compared within and across disease groups. Group comparisons were evaluated for normality using the Shapiro-Wilk test, followed by either the Student’s *t-*test or Wilcoxon rank-sum test, as appropriate. *p*-values and medians with interquartile range were used to annotate module score violin plots.

### CellChat Analysis

The CellChat v2 package was used to assess predicted cell-cell communication, in accordance with their standard workflow (16). For the cell-type-specific CellChat pathway analysis, total signaling strength for each pathway was calculated per cell type by summing outgoing (sender) and incoming (receiver) communication probabilities from the CellChat probability array (@netP$prob). Fold changes in pathway signaling strength were computed for each cell type between disease and control conditions using log2 transformation with a pseudocount of 0.001 to accommodate pathways absent in one condition. Pathways were ranked by absolute fold change magnitude, and the top 5 most altered pathways were selected regardless of statistical significance. Results were visualized as horizontal bar plots displaying log2 fold changes for each disease state relative to their respective controls.

### Human Proteomics Pathway Scores

Proteomic analysis of mitochondrial-enriched fractions from human liver samples was performed as previously described (17). Relative pathway scores were calculated by summing protein abundances of all detectable proteins per sample for each KEGG or WikiPathways pathway. Control samples were averaged, and all samples were then normalized to the control average to generate pathway scores. Error bars represent median with range and *p*-values were calculated using Mann-Whitney tests.

### Rodent Proteomics Pathway Scores

Proteomic analysis of control and monocrotaline-treated rodent livers was performed (17). Pathway scores were calculated using the same approach applied to the human proteomic datasets described above.

### Correlation of HIF-1 Activation with Clinical Markers of RVF/PAH Severity

To assess associations between HIF-1 signaling and clinical severity, we calculated HIF-1 pathway module scores for each cell type using Seurat’s AddModuleScore. Scores were then pseudobulked at the patient level using AggregateExpression, yielding one HIF-1 module score per cell type per patient. These cell type-specific scores were then evaluated for associations with clinical markers of RVF/PAH severity, including hemodynamics (mean pulmonary artery pressure (mPAP), right atrial pressure (RAP), cardiac index, cardiac output, pulmonary vascular resistance (PVR), right ventricular systolic pressure (RVSP), tricuspid annular plane systolic excursion (TAPSE)) and hepatic/systemic biomarkers (ALT, AST, ALP, bilirubin, GGT, NT-proBNP). Pearson correlation coefficients were computed for all variable pairs using pairwise complete observations to accommodate missing clinical data. Correlations were visualized as a hierarchically clustered heatmap (Python seaborn clustermap) and as bar plots showing each cell type-specific HIF-1 score versus individual clinical parameters.

### Figure Preparation

All figures were prepared in Adobe Illustrator 2025.

## Results

### snRNA-seq Identified Specific Alterations in Liver Cellular Composition Among PAH, NASH, and FALD

After processing and removal of low-quality data, we compared 53,057 control nuclei with 71,792 PAH nuclei (Figure 1A). Unsupervised clustering identified eight cell types: hepatocytes, endothelial cells (ECs), hepatic stellate cells (HSCs), macrophages, lymphocytes, B cells, plasma cells, and cholangiocytes. PAH livers had an increased relative abundance of hepatocytes (control: 41.1%, PAH: 60.6%), ECs (control: 9.9%, PAH: 13.4%), plasma cells (control: 0.1%, PAH: 2.0%), and macrophages (control: 4.0%, PAH: 5.5%). However, HSCs (control: 27.7%, PAH: 13.5%), lymphocytes (control: 10.8%, PAH: 3.4%) and cholangiocytes (control: 5.9%, PAH: 1.0%) were reduced.

**Figure 1:**
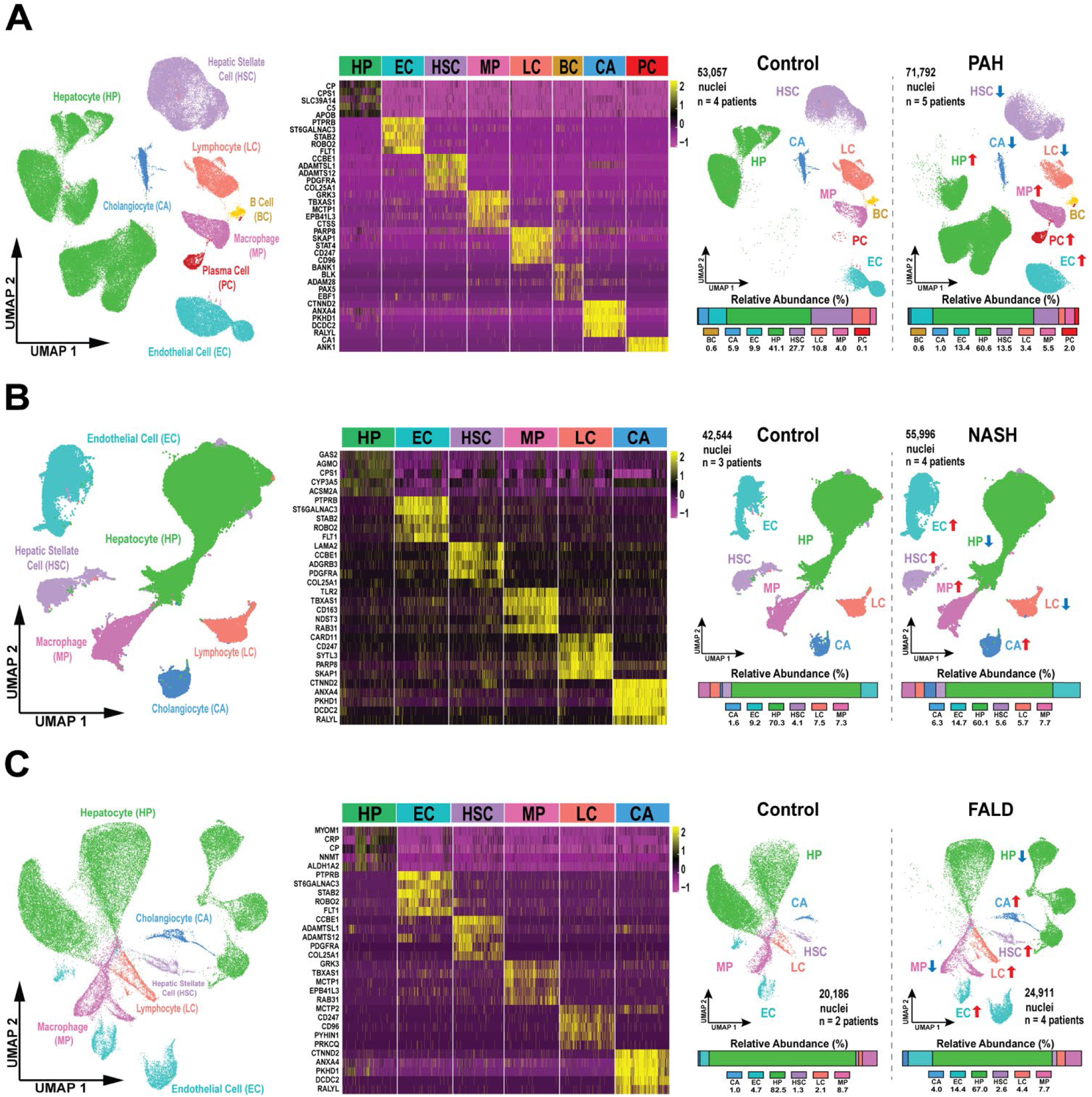
snRNA-seq identified alterations in the cellular landscape in PAH, NASH, and FALD livers. **(A)** Uniform manifold approximation and projection (UMAP) visualization of 8 cell types identified in PAH livers with unsupervised clustering. Validated marker genes used to identify cell type identities in PAH. UMAP and relative abundances of each cell type in control and PAH livers, with arrows indicating relative abundance change in PAH relative to control. **(B)** UMAP visualization of 6 cell types identified in NASH livers with unsupervised clustering. Validated marker genes used to identify cell type identities in NASH. UMAP and relative abundances of each cell type in control and NASH livers, with arrows indicating relative abundance change in NASH relative to control. **(C)** UMAP visualization of 6 cell types identified in FALD livers with unsupervised clustering. Validated marker genes used to identify cell type identities in FALD. UMAP and relative abundances of each cell type in control and FALD livers, with arrows indicating relative abundance change in FALD relative to control.

Next, we compared our results to publicly available data from two distinct triggers of human hepatopathy: NASH (metabolically driven) and FALD (chronic venous stasis driven). The NASH dataset consisted of nuclei from 3 non-NASH controls (42,544 nuclei) and 4 NASH patients (55,996 nuclei) (Figure 1B). Control and NASH livers possessed 6 unique cell types and NASH livers displayed heightened relative abundances of ECs (control: 9.2%, NASH: 14.7%), macrophages (control: 7.3%, NASH: 7.7%), HSCs (control: 4.1%, NASH: 5.6%), and cholangiocytes (control: 1.6%, NASH: 6.3%), but hepatocytes (control: 70.3%, NASH: 60.1%) and lymphocytes (control: 7.5%, NASH: 5.7%) were decreased. We then probed data from 2 non-FALD control (20,186 nuclei) and 4 FALD (24,911 nuclei) patients (Figure 1C). In the FALD dataset, 6 distinct cell types were identified: hepatocytes, ECs, HSCs, macrophages, lymphocytes, and cholangiocytes. FALD livers displayed greater relative abundances of ECs (control: 4.7%, FALD: 13.6%), macrophages (control: 8.3%, FALD: 9.1%), HSCs (control: 1.5%, FALD: 2.5%), cholangiocytes (control: 1.0%, FALD: 3.1%), and lymphocytes (control: 2.0%, FALD: 4.7%), with fewer hepatocytes (control: 82.0%, FALD: 66.9%). Bar plots depicting sample-specific cell-type relative abundances across the three datasets are provided in **Supplemental Figure 3A-C**.

### PAH Hepatocytes Exhibited Metabolic Reprogramming and Altered Cellular Adhesion and Extracellular Matrix Engagement Distinct from NASH and FALD

Unsupervised clustering revealed unique hepatocyte populations in PAH, NASH, and FALD livers relative to their controls (Figure 2A). In PAH hepatocytes, transcripts associated with hypoxia-inducible factor 1 (HIF-1) signaling, carbon metabolism, and glucose metabolism were enriched, whereas those involved in fatty acid metabolism and cytochrome P450 oxidation were reduced (**Supplemental Figure 4A**). NASH hepatocytes enriched fatty acid metabolism transcripts but downregulated cytochrome P450 oxidation (**Supplemental Figure 4B**). FALD hepatocytes demonstrated upregulation of fatty acid and amino acid metabolism, while oxidative phosphorylation transcripts were reduced (**Supplemental Figure 4C**).

**Figure 2:**
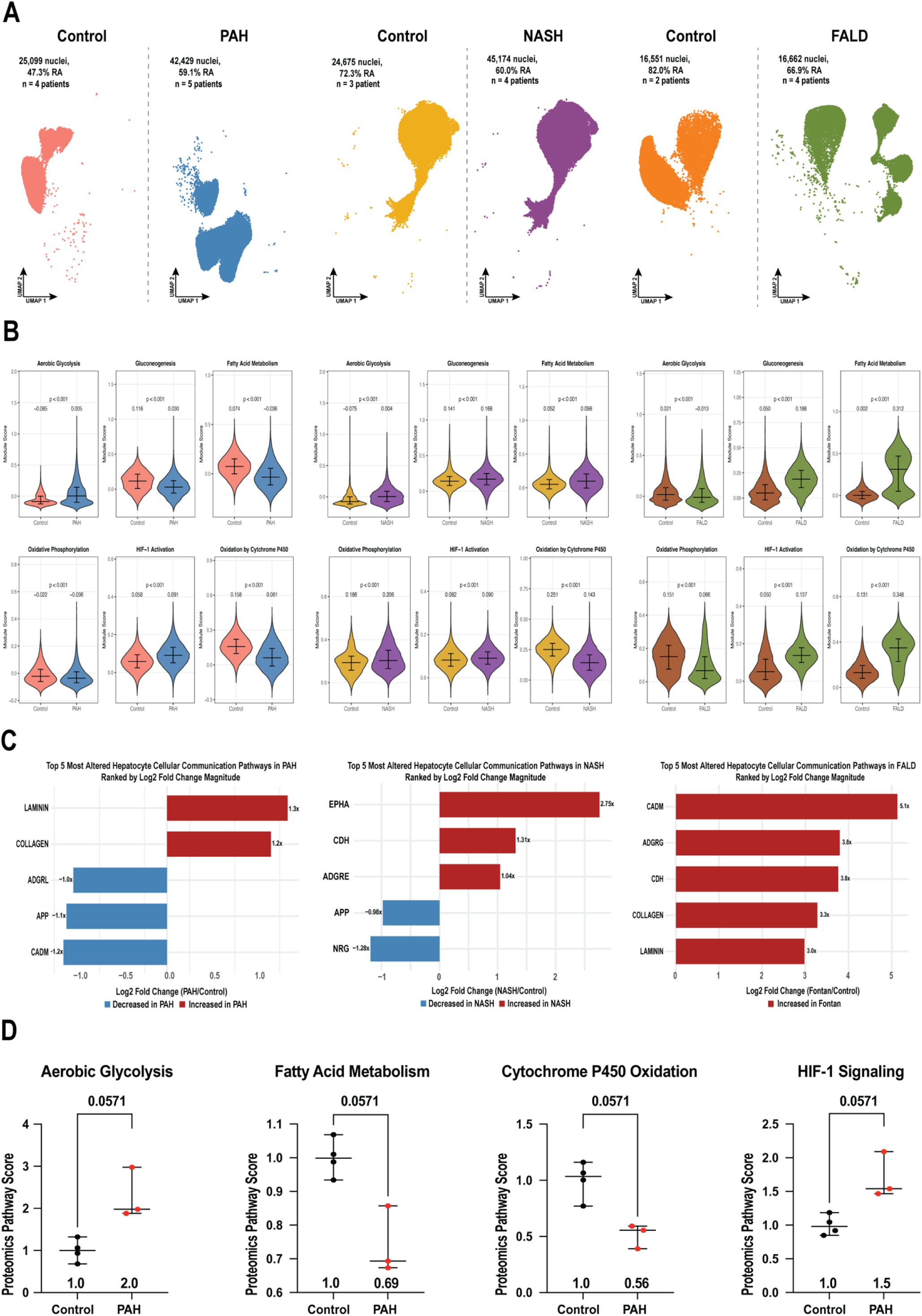
PAH hepatocyte metabolic alterations, pro-proliferative phenotype, and suppressed cell adhesion signaling were distinct from NASH and FALD. **(A)** UMAP visualization of PAH, NASH, and FALD hepatocyte clusters and respective controls. **(B)** Relative pathway activity displayed by violin plots of module scores calculated from relevant enriched/suppressed pathways in PAH, NASH, and FALD hepatocytes. Median values with interquartile range are displayed, and *p*-values were calculated by either Student’s *t-*test or Wilcoxon rank-sum test, as appropriate. **(C)** Top five most dysregulated cellular communication pathways based on total signaling strength across PAH, NASH, and FALD hepatocytes. Cell signaling pathways are ranked based on magnitude of the log2FC change in PAH, with red bars indicating increased pathway signaling in PAH relative to control hepatocytes and blue bars indicating decreased pathways. **(D)** Normalized protein abundances in control and PAH human livers across the WikiPathways aerobic glycolysis pathway and the KEGG fatty acid metabolism, oxidation by cytochrome P450, and HIF-1 signaling pathways. Median values with range are displayed, and *p*-values were calculated using Mann-Whitney tests.

To more precisely probe hepatic metabolism, we calculated module scores to assess engagement in two key metabolic domains: ATP-generating pathways and the cytochrome P450 pathway. In PAH, hepatocytes exhibited heightened aerobic glycolysis and preserved oxidative phosphorylation activity but reduced gluconeogenesis and fatty acid metabolism (Figure 2B). NASH hepatocytes displayed enhanced aerobic glycolysis, gluconeogenesis, fatty acid metabolism, and oxidative phosphorylation activity compared to controls (Figure 2B). Conversely, FALD hepatocytes showed increased fatty acid metabolism, but diminished aerobic glycolysis, gluconeogenesis, and oxidative phosphorylation (Figure 2B). Then, we assessed activity in the cytochrome P450 pathway. Cytochrome P450 activity was suppressed in PAH and NASH, but increased in FALD hepatocytes, indicating disease-specific alterations. Finally, while HIF-1 signaling activity was elevated in all three disease states, the divergent metabolic profiles observed among PAH, NASH, and FALD hepatocytes suggested that HIF-1 activation alone did not account for the full spectrum of metabolic reprogramming (Figure 2B).

We then identified the five most dysregulated signaling pathways in hepatocytes across each disease state using CellChat (16) (Figure 2C). In PAH hepatocytes, ECM signaling via collagen and laminin was elevated. In contrast, cell adhesion molecule, amyloid precursor protein, and adhesion G-protein coupled receptor L signaling was reduced, suggesting suppression of adhesion-related programs (Figure 2C). In NASH hepatocytes, ephrin receptor A signaling, which regulates cell positioning and inflammatory responses (18), was increased alongside cadherin-mediated adhesion and adhesion G-protein-coupled receptor E signaling, which is involved in macrophage-hepatocyte communication. Conversely, signaling through amyloid precursor protein, a mediator of cellular stress responses, and neuregulin, a hepatoprotective growth factor, was reduced (Figure 2C). In FALD, hepatocytes showed signs of structural remodeling, with increased cell adhesion molecule M, adhesion G-protein-coupled receptor G, cadherin, collagen, and laminin signaling (Figure 2C). Interestingly, although hepatocyte remodeling through heightened ECM signaling was present in both PAH and FALD hepatocytes, only FALD displayed activation of adhesion molecules. However, NASH hepatocytes exhibited immune and adhesion-related signaling changes, with less engagement of structural signaling pathways.

Finally, we analyzed proteomic data from mitochondrial-enriched fractions of a subset of the same control and PAH livers (*n*=4 control, *n*=3 PAH) to determine whether these metabolic alterations were present at the protein level. Consistent with our transcriptomic findings, normalized protein abundances in the aerobic glycolysis and HIF-1 signaling pathways were non-significantly increased in PAH livers, whereas proteins involved in the fatty acid metabolism and cytochrome P450 oxidation pathways were non-significantly reduced relative to controls (Figure 2D).

### PAH Endothelial Cells Demonstrated Glycolytic Bias and Altered Barrier Function

Next, we evaluated EC reprogramming in PAH, NASH, and FALD livers (Figure 3A). In PAH, ECs demonstrated hypoxia-associated metabolic remodeling as HIF-1 signaling, glycolysis, PI3K-Akt, and pentose phosphate pathways were enriched (**Supplemental Figure 5A**). In contrast, transcripts related to the innate immune system, type I interferon signaling, and FoxO signaling were suppressed (**Supplemental Figure 5A**). NASH ECs showed heightened ribosome, fatty acid metabolism, and complement pathway transcripts but downregulation of one carbon metabolism and AMPK signaling (**Supplemental Figure 5B**). FALD ECs upregulated Ras, focal adhesion, and B cell receptor signaling, while oxidative phosphorylation and complement signaling transcripts were reduced (**Supplemental Figure 5C**).

**Figure 3:**
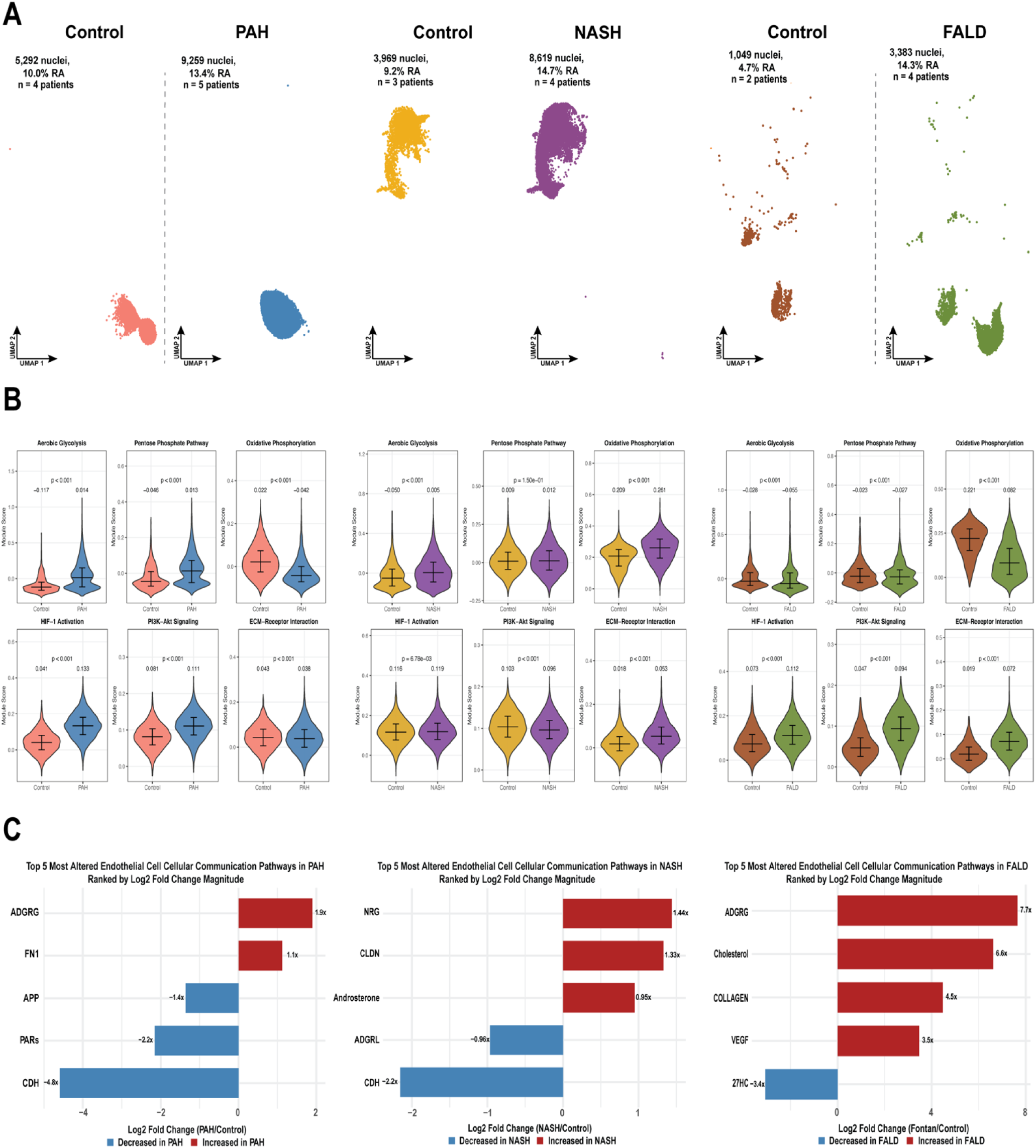
PAH endothelial cells exhibited Warburg-like metabolism and alterations in cell adhesion and barrier function pathways. **(A)** UMAP visualization of PAH, NASH, and FALD endothelial cell clusters and respective controls. **(B)** Relative pathway activity displayed by violin plots of module scores calculated from relevant enriched/suppressed pathways in PAH, NASH, and FALD endothelial cells. Median values with interquartile range are displayed, and *p*-values were calculated by either Student’s *t-*test or Wilcoxon rank-sum test, as appropriate. **(C)** Top five most dysregulated cellular communication pathways based on total signaling strength across PAH, NASH, and FALD endothelial cells. Cell signaling pathways are ranked based on magnitude of the log2FC change in PAH, with red bars indicating increased pathway signaling in PAH relative to control endothelial cells and blue bars indicating decreased pathways.

We then calculated module scores to assess engagement in core glycolytic and growth/survival signaling pathways across the disease states (Figure 3B). In PAH, ECs showed elevated activity in the aerobic glycolysis and pentose phosphate pathways but reduced oxidative phosphorylation. NASH ECs displayed increased aerobic glycolysis and oxidative phosphorylation with preserved pentose phosphate pathway activity. In contrast, FALD ECs had comparable aerobic glycolysis and pentose phosphate activity relative to controls but suppressed oxidative phosphorylation (Figure 3B). We next assessed PI3K-Akt and ECM-receptor interaction activity. PAH ECs demonstrated increased PI3K-Akt signaling with minimally altered ECM-receptor interaction activity relative to control ECs. NASH ECs augmented ECM-receptor interaction activity and preserved PI3K-Akt signaling while FALD ECs upregulated both pathways (Figure 3B).

To probe how these cellular changes were predicted to modulate cell-cell interactions, we identified the five most dysregulated EC cellular communication pathways in each disease state. In PAH ECs, cell adhesion signaling via adhesion G-protein-coupled receptor G and fibronectin-1 was increased, while signaling related to cadherin, protease-activated receptor and amyloid precursor protein was reduced (Figure 3C), suggesting EC barrier function may be compromised as both cadherin and protease-activated receptor are central to endothelial barrier integrity and vascular permeability (19,20). In NASH ECs, neuregulin, which promotes vascular homeostasis, was elevated along with claudin, a regulator of tight junction integrity (21), and androsterone signaling. In contrast, signaling via adhesion G-protein-coupled receptor L and cadherins was reduced (Figure 3C). In FALD, ECs displayed increased matrix and angiogenic signaling, including elevated adhesion G-protein-coupled receptor G, cholesterol, collagen, and VEGF signaling, while 27-hydroxycholesterol signaling was reduced (Figure 3C). Thus, EC signaling appeared to be distinct across diseases. PAH had alterations in cell adhesion and barrier function; FALD ECs exhibited structural and angiogenic remodeling, and NASH ECs showed heightened steroid signaling with dysregulation of vascular homeostasis pathways.

### Divergent Metabolic and Fibrotic Signaling Profiles Distinguished Hepatic Stellate Cells in PAH, NASH, and FALD

We next examined transcriptional alterations in HSCs (Figure 4A). In PAH, HSCs demonstrated increased HIF-1 and PI3K-Akt transcripts, with concurrent enrichment of carbon, glucose, and amino acid metabolism. Conversely, transcripts involved in innate immune sensing, transforming growth factor beta (TGF-β), and fatty acid metabolism were suppressed (**Supplemental Figure 6A**). NASH HSCs exhibited upregulation of ECM-receptor interaction, PI3K-Akt, and focal adhesion transcripts, accompanied by downregulation of Wnt signaling and dysregulated metabolism (**Supplemental Figure 6B**). In FALD, HSCs displayed enhancement of TGF-β and Wnt signaling transcripts while oxidative phosphorylation was reduced (**Supplemental Figure 6C**).

**Figure 4:**
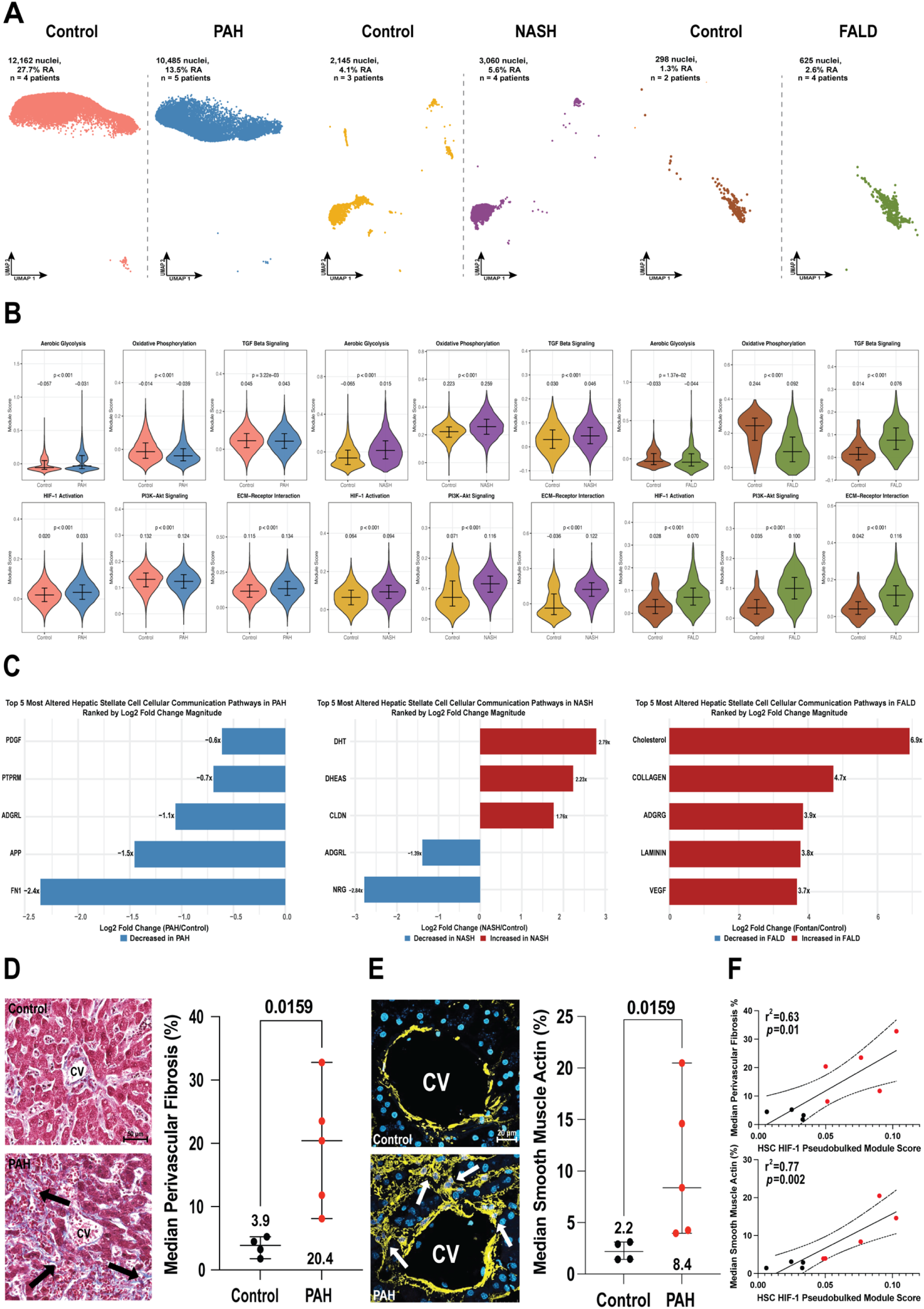
Upregulated HIF-1 and PI3K-Akt signaling characterized hepatic stellate cell signaling in PAH. **(A)** UMAP visualization of PAH, NASH, and FALD hepatic stellate cell clusters and respective controls. **(B)** Relative pathway activity displayed by violin plots of module scores calculated from relevant enriched/suppressed pathways in PAH, NASH, and FALD hepatic stellate cells. Median values with interquartile range are displayed, and *p*-values were calculated by either Student’s *t-*test or Wilcoxon rank-sum test, as appropriate. **(C)** Top five most dysregulated cellular communication pathways based on total signaling strength across PAH, NASH, and FALD hepatic stellate cells. Cell signaling pathways are ranked based on magnitude of the log2FC change in PAH, with red bars indicating increased pathway signaling in PAH relative to control hepatic stellate cells and blue bars indicating decreased pathways. **(D)** Representative images and quantification of median perivascular fibrosis (%) surrounding the central vein (CV) in control and PAH livers using Trichrome staining (blue indicates fibrosis). Error bars represent median values with range, and *p*-values were calculated using Mann–Whitney tests. Arrows indicate peri-vascular fibrosis. **(E)** Representative images and quantification of median α–smooth muscle actin (%) surrounding the central vein in control and PAH livers. Yellow indicates α–smooth muscle actin staining and blue indicates DAPI-stained nuclei. Error bars represent median values with range, and *p*-values were calculated using Mann–Whitney tests. Arrows highlight alpha-smooth positive HSCs. **(F)** Linear regression analyses showing associations between HSC HIF-1 activation and median perivascular fibrosis (top) and mean α–smooth muscle actin (bottom) in control (black) and PAH (red) patients. R² and *p*-values were calculated using linear regression. Dotted lines indicate 95% confidence intervals.

We then evaluated metabolic and pro-fibrotic pathways in HSCs from PAH, NASH, and FALD livers (Figure 4B). There were divergent metabolic alterations with regards to glycolysis and oxidative phosphorylation. NASH HSCs exhibited heightened metabolic regulation with increased glycolysis and oxidative phosphorylation while these pathways were minimally altered or suppressed in PAH and FALD (Figure 4B). We then assessed PI3K-Akt, ECM-receptor interaction, and TGF-β signaling due to their roles in hepatic fibrosis. ECM-receptor interaction transcript expression was increased in all three disease states, but PI3K-Akt and TGF-β pathways were reduced in PAH despite being elevated in both NASH and FALD (Figure 4B). Interestingly, HIF-1 activity, which regulates fibrogenic activation under hypoxic stress, was consistently elevated in HSCs across disease states (Figure 4B).

Next, we identified the five most altered HSC cellular communication pathways in each disease state (Figure 4C). In PAH, HSCs showed reduced signaling via several cell adhesion molecules and growth factors, including fibronectin-1, amyloid precursor protein, adhesion G-protein-coupled receptor L, receptor-type tyrosine-protein phosphatase M, and platelet-derived growth factor (Figure 4C). NASH HSCs displayed enriched inflammation-related steroid hormone signaling, evidenced by elevated dihydrotestosterone and dehydroepiandrosterone sulfate activity, as well as increased claudin activity while adhesion G-protein-coupled receptor L and neuregulin signaling was suppressed (Figure 4C). In FALD, HSCs had heightened structural remodeling and pro-fibrotic communication via increased collagen, adhesion G-protein-coupled receptor G, laminin, and vascular endothelial growth factor (VEGF) signaling (Figure 4C). Together, these data demonstrate distinct, disease-specific reprogramming of cellular signaling in HSCs from PAH, NASH, and FALD livers.

Finally, we examined whether transcriptomic changes in PAH HSCs were accompanied by histologic evidence of HSC activation and fibrotic remodeling. PAH livers displayed increased total fibrosis relative to controls, although this change did not reach statistical significance (**Supplemental Figure 7A**). However, sample-specific HIF-1 signaling in HSCs was positively and significantly correlated with total fibrosis in liver specimens (**Supplemental Figure 7B**). When we examined perivascular fibrosis surrounding hepatic central veins, we found a significant elevation in PAH livers (Figure 4D). In addition, alpha-smooth muscle actin immunoreactivity, an indicator of activated HSCs (22), was also significantly heightened in PAH livers compared to controls (Figure 4E). Both median perivascular fibrosis and mean alpha-smooth muscle actin positive area were positively and significantly associated with sample-specific HIF-1 signaling in HSCs (Figure 4F).

### PAH Macrophages Exhibited Complement Activation, Oxidative Metabolism, and Immune Signaling that are Distinct from NASH and FALD Macrophages

We then examined transcriptional changes in hepatic macrophages from PAH, NASH, and FALD livers (Figure 5A). In PAH, macrophages upregulated complement and coagulation transcripts, while JAK-STAT signaling and fatty acid metabolism were suppressed (**Supplemental Figure 8A**). NASH macrophages showed elevated transcripts in the complement, ferroptosis, ribosome pathways, but downregulation of PDGFR-beta and Wnt signaling (**Supplemental Figure 8B**). In FALD, macrophages exhibited increased platelet activation, IL-3, and chemokine signaling transcripts, alongside diminished oxidative phosphorylation (**Supplemental Figure 8C**).

**Figure 5:**
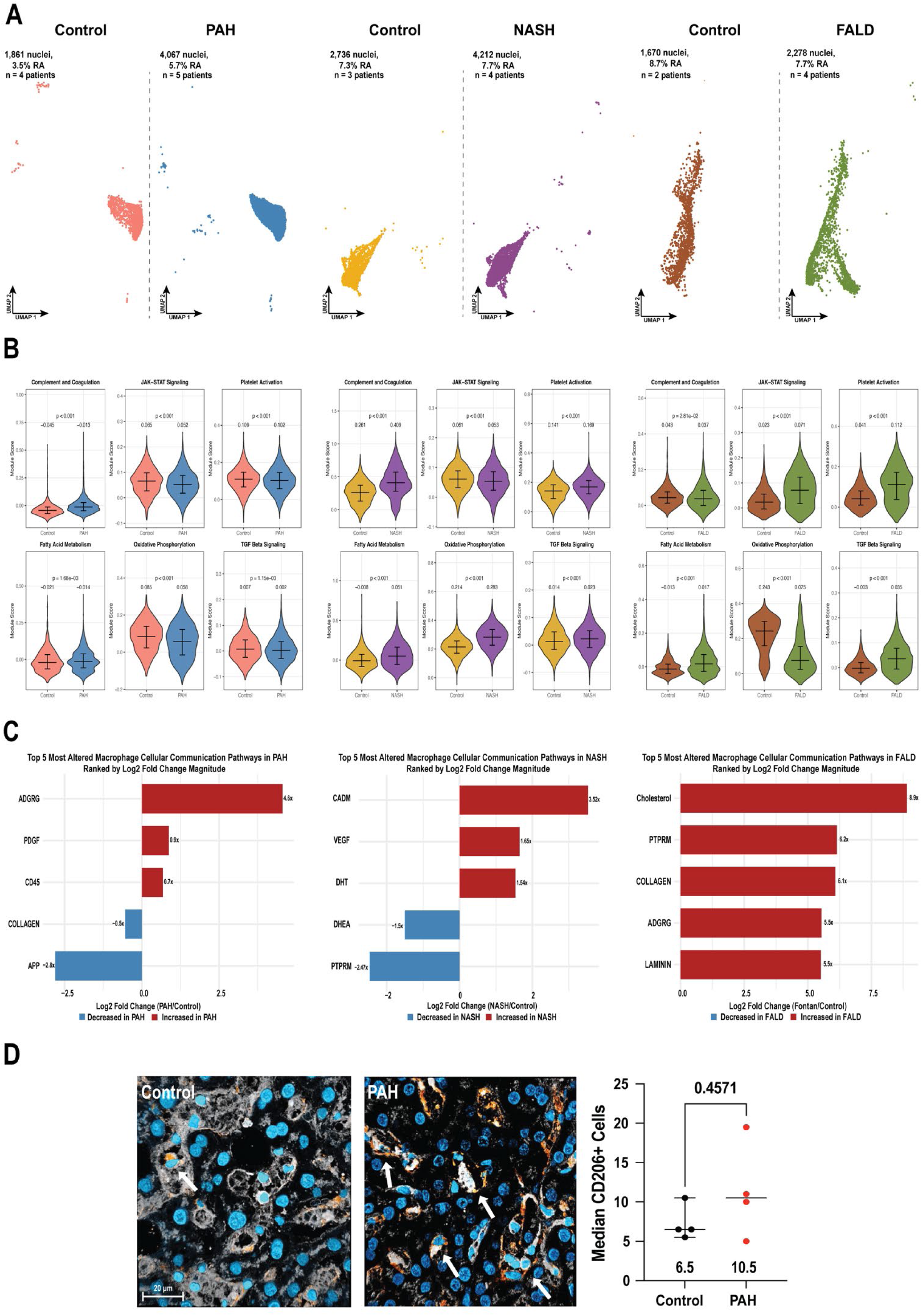
Pro-inflammatory complement signaling with suppressed JAK-STAT signaling and cell-cell communication defined PAH macrophages. **(A)** UMAP visualization of PAH, NASH, and FALD macrophage clusters and respective controls. **(B)** Relative pathway activity displayed by violin plots of module scores calculated from relevant enriched/suppressed pathways in PAH, NASH, and FALD macrophages. Median values with interquartile range are displayed, and *p-*values were calculated by either Student’s t-test or Wilcoxon rank-sum test, as appropriate. **(C)** Top five most dysregulated cellular communication pathways based on total signaling strength across PAH, NASH, and FALD macrophages. Cell signaling pathways are ranked based on magnitude of the log2FC change in PAH, with red bars indicating increased pathway signaling in PAH relative to control macrophages and blue bars indicating decreased pathways. **(D)** Representative images and quantification of median CD206⁺ cells per area in control and PAH livers. Orange indicates CD206 staining, gray indicates WGA, and blue indicates DAPI-stained nuclei. Error bars represent median values with range, and *p*-values were calculated using Mann–Whitney tests.

Next, we assessed whether hepatic macrophages from PAH, NASH, and FALD livers demonstrated disease-specific regulation of immune, metabolic, and fibrogenic pathways (Figure 5B). PAH and NASH macrophages had elevated complement activity and reduced JAK-STAT signaling, while FALD macrophages maintained complement activity and upregulated JAK-STAT signaling (Figure 5B). Metabolically, PAH macrophages retained fatty acid metabolism activity but showed diminished oxidative phosphorylation (Figure 5B). NASH macrophages had increased activity in both pathways, whereas FALD macrophages displayed elevated fatty acid metabolism but suppressed oxidative phosphorylation (Figure 5B). Then, analysis of the TGF-β pathway revealed minimally altered TGF-β activity in PAH but increased signaling in NASH and FALD macrophages (Figure 5B).

We then identified the five most dysregulated macrophage cellular communication pathways in PAH, NASH, and FALD livers (Figure 5C). In PAH, macrophages exhibited enhanced signaling in pathways related to leukocyte migration (23) and immune activation (adhesion G-protein-coupled receptor G, platelet-derived growth factor, and CD45) (24,25), while amyloid precursor protein and collagen signaling was reduced (Figure 5C). NASH macrophages showed upregulation of cell adhesion molecule signaling as well as elevated VEGF and dihydrotestosterone signaling, both of which are implicated in angiogenesis and immune activation. In contrast, signaling through dehydroepiandrosterone, an anti-inflammatory steroid precursor, and protein tyrosine receptor type M was suppressed (Figure 5C). In FALD, macrophages displayed elevated cholesterol signaling, along with enrichment of receptor-type tyrosine-protein phosphatase M, collagen, adhesion G-protein-coupled receptor G, and laminin signaling (Figure 5C). In summary, these data suggest that PAH macrophage signaling is marked by immune activation, NASH by pro-inflammatory steroid hormone signaling, and FALD by pro-fibrotic and metabolic cues.

Finally, we assessed whether the increased macrophage relative abundances observed transcriptionally in PAH livers were reflected histologically. Immunohistochemical staining for CD206 (MRC1), a well-established macrophage marker (26), demonstrated heightened CD206+ cells in PAH livers compared with controls (Figure 5D), but the difference did not reach statistical significance.

### Potential Systemic Implications of PAH Hepatopathy

Given the liver’s role in regulating systemic physiology through secretion of vasoactive peptides, inflammatory cytokines, and metabolic messengers (27,28), we analyzed changes in each of these respective pathways. Fms-related receptor tyrosine kinase 1 (FLT1), a key VEGF receptor involved in angiogenesis (29), was significantly upregulated in PAH hepatocytes and ECs (Figure 6A). GDF15, a pro-proliferative cytokine (30), was also increased in PAH hepatocytes. PAH HSCs had elevated expression of ENG, which encodes vasoconstrictive protein endoglin (31). In contrast, GDF2, the precursor to the vasodilatory protein BMP9 (32), was decreased in PAH hepatocytes and HSCs (Figure 6A). Interleukin-6 (IL-6), a prognostic biomarker in PAH (33), was the most differentially regulated transcript in PAH HSCs. Elevated IL-6 expression was observed in PAH HSCs, macrophages, and hepatocytes, but statistical significance was only reached in HSCs (Figure 6B). Finally, we evaluated ketone metabolism in PAH livers and found the ketone pathway was significantly reduced in PAH hepatocytes (Figure 6C). Collectively, these findings suggested that PAH hepatopathy may exacerbate pulmonary vascular disease and other systemic processes through dysregulated vasoactive signaling, heightened expression of the inflammatory cytokine IL-6, and suppression of ketone body synthesis.

**Figure 6:**
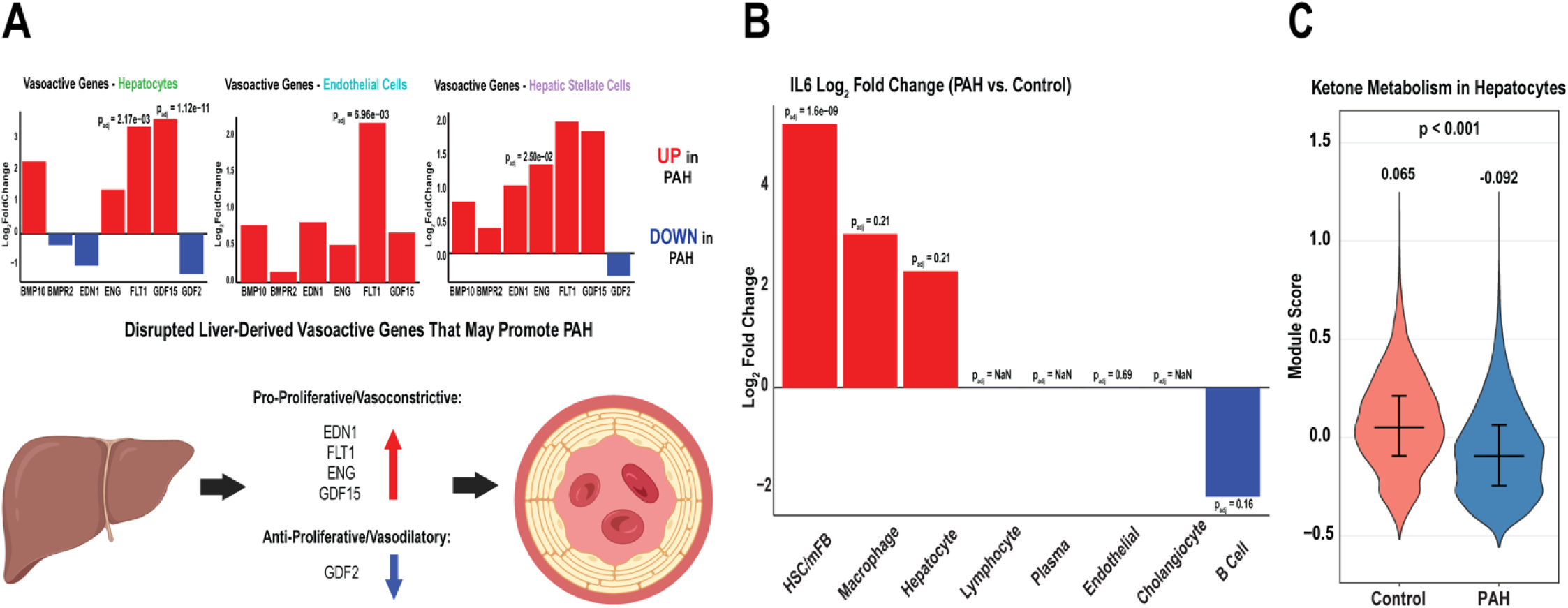
PAH altered the expression of vasoactive peptides in multiple cell types, heightened interleukin-6 production in hepatic stellate cells, and suppressed ketone metabolism in hepatocytes. **(A)** Relative log2FC change in expression between control and PAH for relevant pro-proliferative and vasoactive genes across hepatocytes, endothelial cells, and hepatic stellate cells. *p*-values were calculated using a Wilcoxon rank-sum test. Graphical representation of predicted consequences stemming from dysregulated expression of selected vasoactive genes in the liver on pulmonary vascular remodeling in PAH. **(B)** Relative log2FC change in expression between control and PAH for IL-6 production across all liver cell types. *p*-values were calculated using a Wilcoxon rank-sum test. **(C)** Relative ketone metabolism activity displayed by a violin plot of the hepatocyte module scores for the ketone metabolism pathway. Significance was assessed via a Wilcoxon rank-sum test.

## Discussion

Here, we present a comprehensive single-cell transcriptomic atlas of PAH hepatopathy that identifies cell-specific alterations in liver composition, metabolism, and intercellular signaling. By comparing PAH with NASH and FALD, we define key molecular features that distinguish PAH hepatopathy from other human liver diseases with metabolic or venous stasis triggers. Our snRNA-seq analysis demonstrates that PAH most prominently alters hepatocyte metabolism, promoting a pro-proliferative, hypoxia-adapted, Warburg-like (34) metabolic phenotype. In contrast, NASH hepatocytes exhibit a hypermetabolic state, whereas FALD hepatocytes selectively suppress oxidative phosphorylation without enhancing glycolysis. Transcripts associated with the cytochrome P450 pathway decrease in PAH and NASH hepatocytes but increase in FALD. Although hepatocytes from all three diseases activate extracellular matrix (ECM) signaling programs, only PAH broadly suppresses cell adhesion pathways. Notably, whole-liver proteomic data from a subset of the same control and PAH patients recapitulate the hepatocyte metabolic alterations observed in our transcriptomic analysis, providing orthogonal support for these findings.

Our comparative analysis suggests PAH, NASH, and FALD have cell-type specific alterations that distinguish the three diseased states. In hepatic stellate cells (HSCs), PAH and NASH enrich PI3K–Akt signaling transcripts, whereas NASH and FALD co-activate PI3K–Akt and TGF-β signaling pathways. Consistent with these transcriptional changes, PAH livers exhibit prominent perivascular fibrosis with increased α-smooth muscle actin immunoreactivity indicating a greater abundance of activated HSCs surrounding the central vein. Macrophage programs also differ across diseases: PAH and NASH macrophages both induce pro-inflammatory complement signaling, but only PAH macrophages show reduced predicted cell–cell interactions alongside amplified immune signaling. In contrast, FALD macrophages downregulate complement signaling but uniquely upregulate JAK–STAT signaling. Across multiple liver cell types, PAH also modulates the expression of vasoactive genes, increases IL-6 production in HSCs, and suppresses the hepatocyte ketone metabolism program. Together, these findings implicate the liver as an active contributor to PAH pathobiology and identify disease-specific pathways that may be targeted to improve hepatic function.

To begin proposing a mechanistic model of how PAH alters hepatic function, we examined whether HIF-1 activity in hepatocytes, ECs, and HSCs correlates with clinical markers of PAH severity. HIF-1 signaling in all three cell types positively correlates with mean pulmonary artery pressure (mPAP), pulmonary vascular resistance (PVR), and right atrial pressure (RAP) (**Supplemental Figure 9A**). Among these populations, HIF-1 activation in HSCs, the principal drivers of hepatic fibrosis (35), shows the strongest association with RVF/PAH severity and significantly correlates with mPAP (**Supplemental Figure 9B**). Therefore, we explored a potential molecular mechanism linking hemodynamic disturbances to altered transcriptional programs in HSCs and hepatocytes. Because hepatic congestion and regurgitant flow resulting from tricuspid regurgitation may impose mechanical stress on the liver, we examined the expression of Piezo mechanosensitive ion channels (36) in HSCs. Piezo1, but not Piezo2, is significantly elevated in PAH HSCs and correlates with perivascular fibrosis (**Supplemental Figure 10A**). As increased Piezo1 activity promotes HIF-1 activation and IL-6 secretion in other cell types (37,38), we assessed relationships among Piezo1, HIF-1, and IL-6 in HSCs. Indeed, HSC Piezo1 expression positively correlates with both HIF-1 activity and IL-6 expression (**Supplemental Figure 10B**). These findings suggest a potential pathway whereby hemodynamic stress increases Piezo1 signaling in HSCs, triggering HIF-1 activation and IL-6 production. Because IL-6 can induce HIF-1 activation in hepatocytes (39), we hypothesize that HSC-derived IL-6 acts as a paracrine signal to promote the Warburg-like metabolic reprogramming observed in PAH hepatocytes. In summary, this working model links altered hemodynamics in PAH to Piezo1-dependent activation of HSCs, which then restructures hepatocyte metabolism through IL-6–mediated HIF-1 signaling (**Supplemental Figure 10C**). Although this hypothesis is supported by our correlative analyses, experimental validation will undoubtedly be required to establish causality.

We find PAH hepatocytes exhibit increased HIF-1 signaling, which may have important systemic metabolic consequences. Hypoxia and HIF-1 can suppress cytochrome P450–mediated oxidation (40), potentially impairing drug metabolism, lipid processing, bile acid synthesis, and detoxification of reactive metabolites. Reduced cytochrome P450 activity may also contribute to PAH-associated dyslipidemia (41) by limiting cholesterol hydroxylation and fatty acid oxidation. In addition, HIF-1 activation may disrupt ketone body synthesis, as constitutive HIF-1 signaling suppresses ketogenesis in experimental models (42). Because ketone bodies are an energetically efficient fuel source with anti-inflammatory properties (28), impaired ketone production could exacerbate systemic energy deficits and promote inflammatory signaling. This mechanism may help explain why compensatory ketosis is often absent in PAH patients with severe right ventricular dysfunction (43). Modulating hepatocyte HIF-1 signaling may therefore represent a potential therapeutic strategy to restore drug metabolism, improve ketone production, and alleviate systemic metabolic dysfunction in PAH.

Endothelial dysfunction in the pulmonary vasculature defines PAH (44), and our data suggest that this phenomenon extends to the liver. PAH ECs adopt a hypoxia-adapted, Warburg-like metabolic phenotype (34), characterized by increased glycolysis and HIF-1 signaling, and EC HIF-1 activation displays the second strongest correlation with markers of RVF/PAH severity after HSCs. Because liver sinusoidal ECs line the hepatic sinusoids and are highly responsive to shear stress (45), congestion-induced disturbances in sinusoidal flow may also promote EC HIF-1 activation and metabolic reprogramming, paralleling the responses observed in HSCs. Again, this hypothesis will require additional validation.

Finally, our data suggest PAH macrophages upregulate complement signaling. Locally, complement deposition can damage hepatocytes and other liver cells by triggering innate immune responses that lead to fibrosis and scarring (46). Although selective suppression of JAK/STAT signaling may dampen pro-inflammatory cytokine secretion and attenuate fibrogenesis, reduced STAT3 activity could be deleterious as it may also impair efferocytosis-mediated liver regeneration (47). Therefore, downregulation of JAK/STAT signaling in PAH macrophages may limit fibrosis but impair efferocytosis and regeneration.

### Limitations

Our study has several important limitations. First, nuclei isolation methods differed across the PAH, NASH, and FALD datasets; to improve comparability, we reanalyzed raw NASH and FALD data using the same Seurat pipeline as the PAH dataset. Second, as a single-nucleus RNA sequencing study, our data lack spatial resolution and cannot capture zonal heterogeneity within the hepatic lobule, highlighting the need for future spatial transcriptomic approaches. Third, the modest sample size inherent to rare human tissue studies limits statistical power and increases susceptibility to type I and type II error, necessitating cautious interpretation of association-based findings. Lastly, the PAH and NASH samples were autopsy-derived and therefore likely reflect end-stage disease, which may limit generalizability to earlier disease stages. Consistent with this interpretation, mitochondrial proteomic analysis of monocrotaline-treated rodent livers demonstrated similar reductions in fatty acid metabolism and oxidative phosphorylation, supporting cross-species metabolic convergence (**Supplemental Figure 11**). However, HIF-1 signaling was not increased in the rodents, which may reflect differences in disease chronicity or end-stage disease.

## Acknowledgements

We are grateful to Shuang Wang and Po Hu for their assistance in obtaining the NASH and FALD datasets, respectively. We thank the University of Minnesota Genomics Center, who performed the sequencing for the single-nucleus RNA sequencing experiment in this manuscript.

## Central Figure/Graphical Abstract

**Supplemental Figure 1:** Schematic representation of single nucleus RNA-sequencing workflow and data processing pipeline.

**Supplemental Figure 2:** Violin plots displaying key snRNA-seq quality control metrics across datasets, including total reads per nuclei, detected genes per cell, percent mitochondrial reads, and percent ribosomal reads. Median values are displayed and error bars represent medians with interquartile range.

**Supplemental Figure 3:**
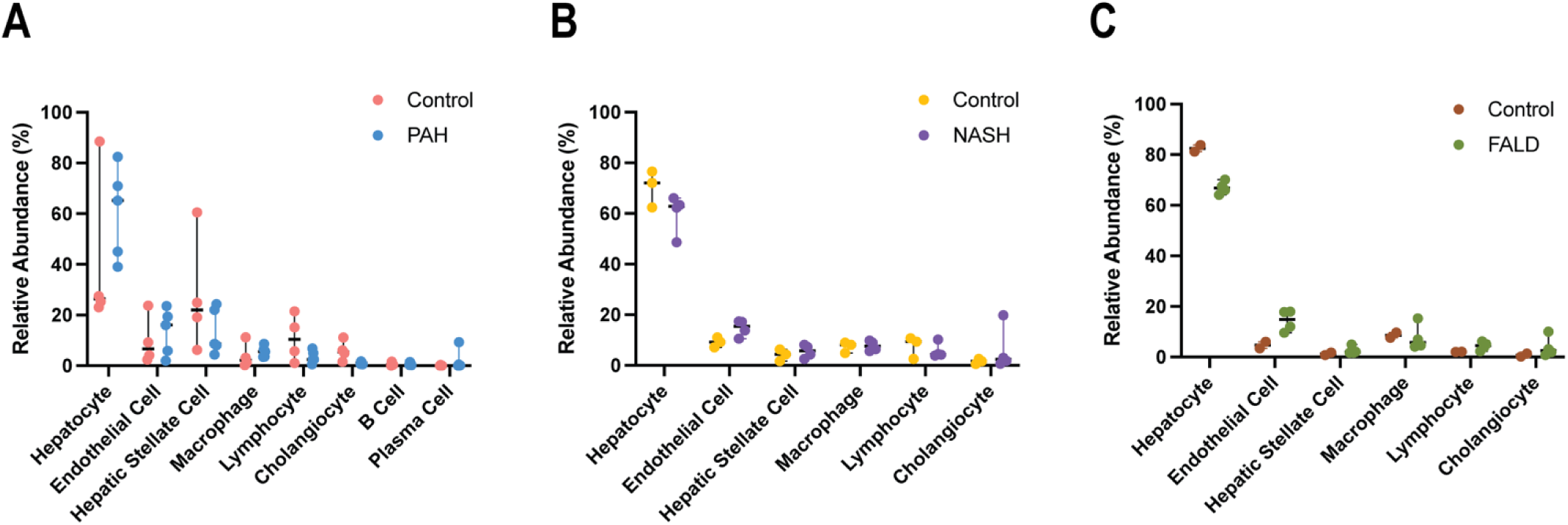
Sample-specific relative abundance values across cell types in PAH, NASH, and FALD datasets. (A) Cell-type relative abundances in control and PAH samples. Error bars represent median values with range. (B) Cell-type relative abundances in control and NASH samples. Error bars represent median values with range. (C) Cell-type relative abundances in control and FALD samples. Error bars represent median values with range.

**Supplemental Figure 4:**
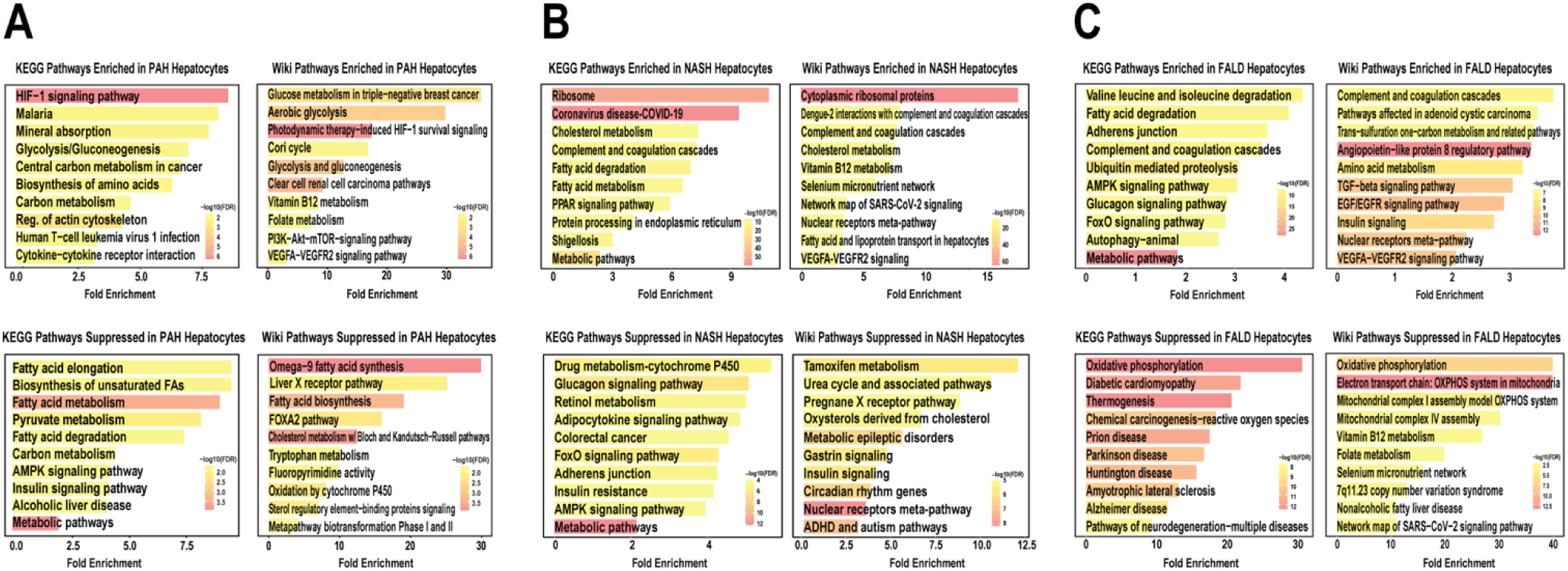
KEGG and Wiki pathway analysis of enriched and suppressed DEGs from PAH, NASH, and FALD hepatocytes relative to their respective controls.

**Supplemental Figure 5:**
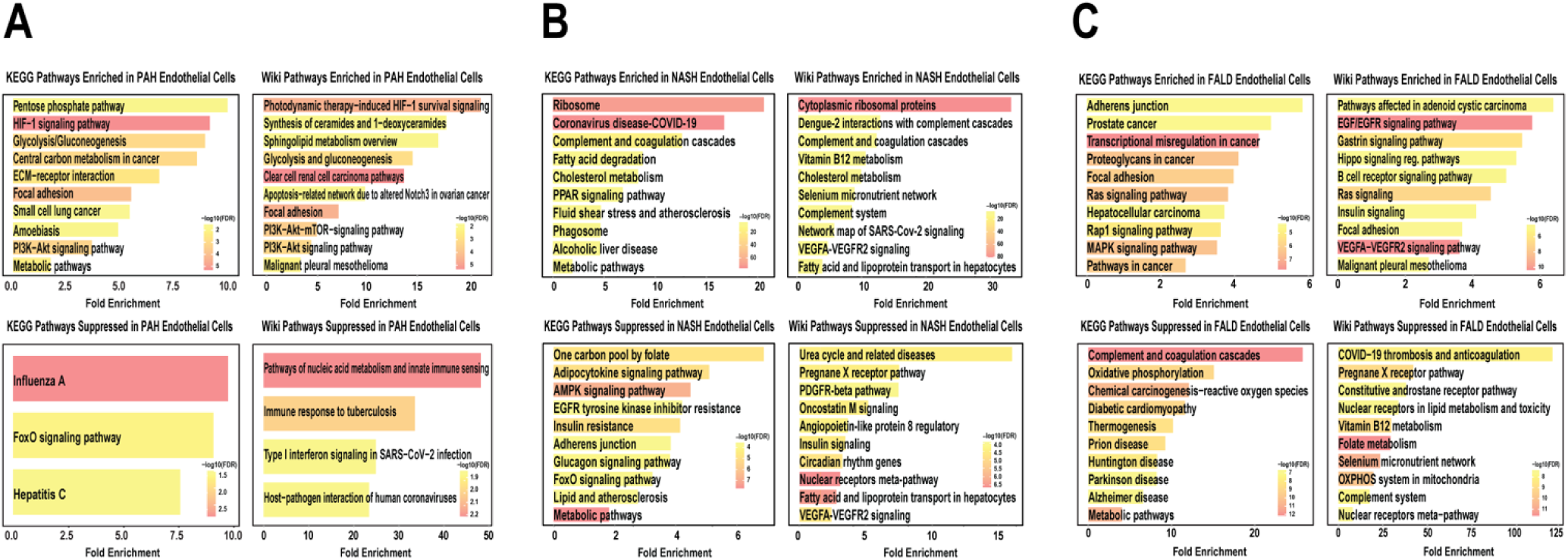
KEGG and Wiki pathway analysis of enriched and suppressed DEGs from PAH, NASH, and FALD endothelial cells relative to their respective controls.

**Supplemental Figure 6:**
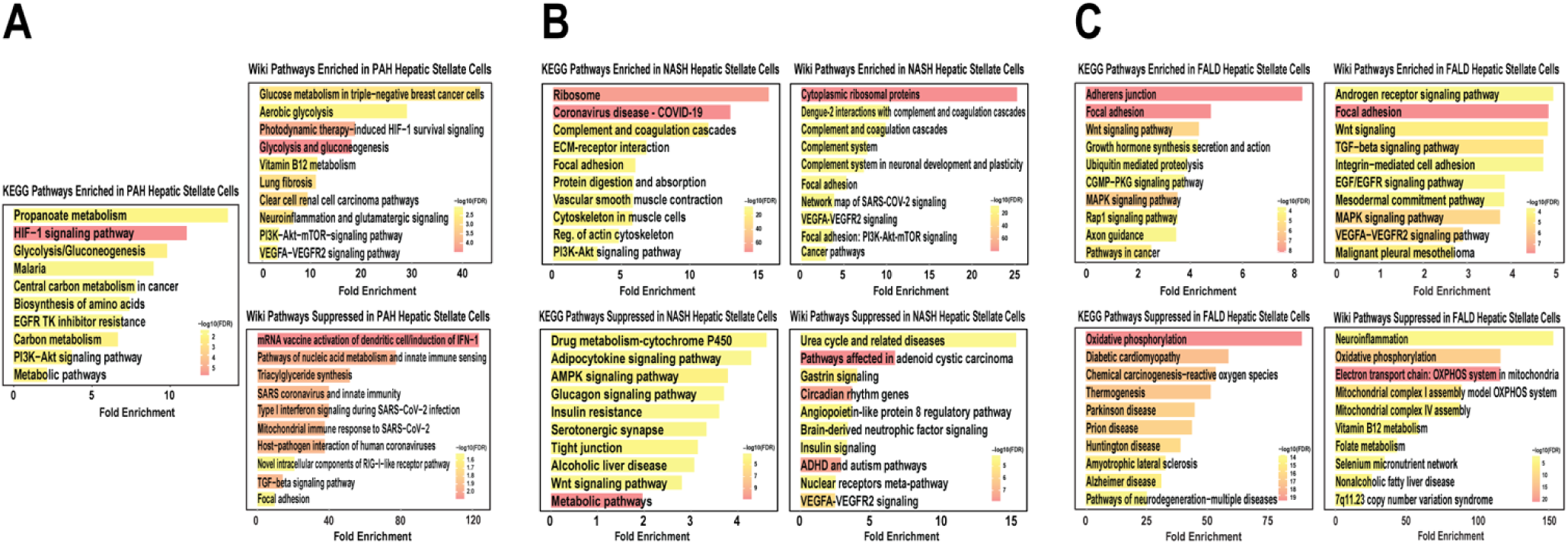
KEGG and Wiki pathway analysis of enriched and suppressed DEGs from PAH, NASH, and FALD hepatic stellate cells relative to their respective controls.

**Supplemental Figure 7:**
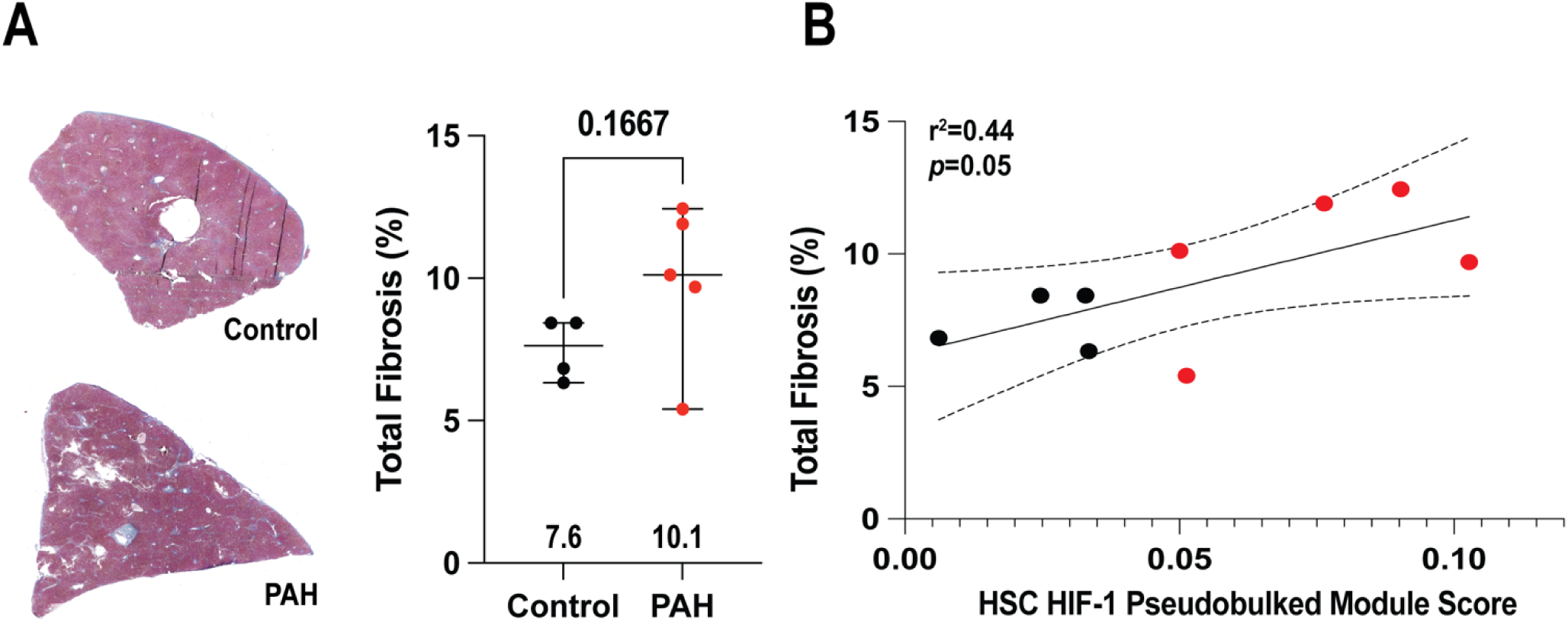
Total hepatic fibrosis was elevated in PAH livers and significantly associated with hepatic stellate cell HIF-1 signaling. (A) Representative whole-liver images and quantification of trichrome total fibrosis (%) staining in control and PAH livers. Error bars represent median values with range, and the *p*-value was calculated by Mann-Whitney test. (B) Linear regression analysis showing the association between total fibrosis (%) and hepatic stellate cell HIF-1 pseudobulked module scores in control (black) and PAH (red) patients. Dotted lines display 95% confidence intervals.

**Supplemental Figure 8:**
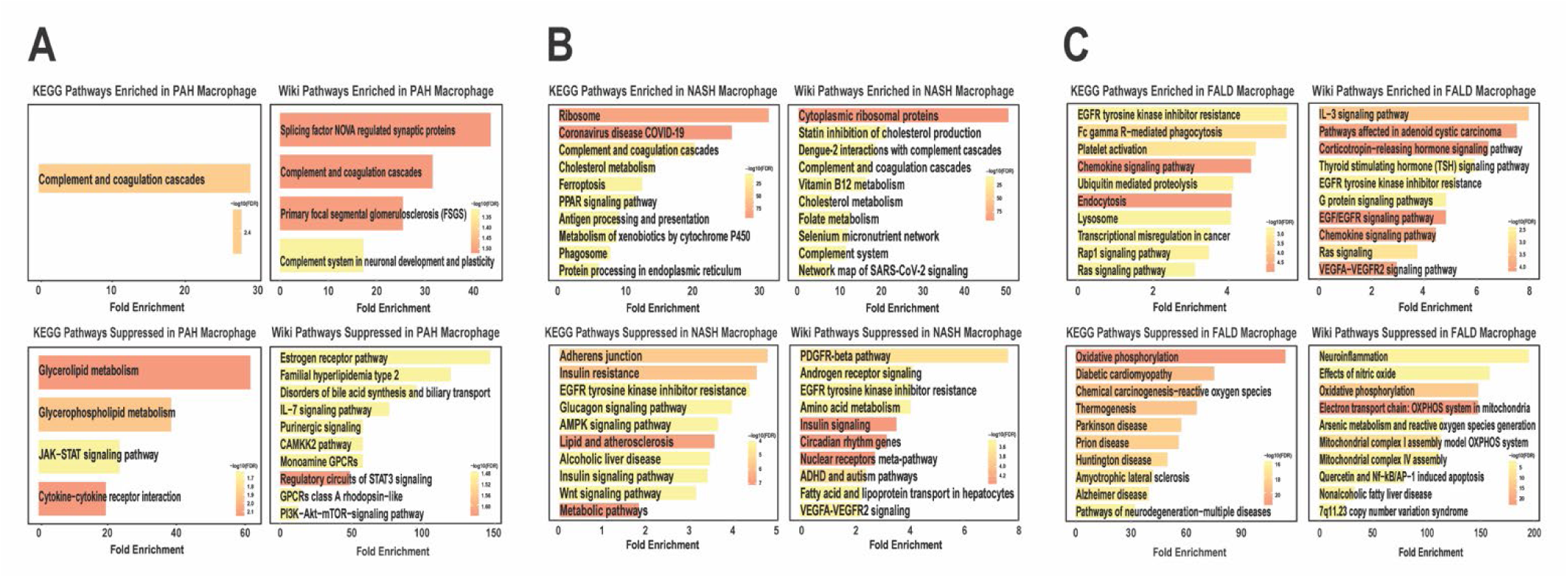
KEGG and Wiki pathway analysis of enriched and suppressed DEGs from PAH, NASH, and FALD macrophages relative to their respective controls.

**Supplemental Figure 9:**
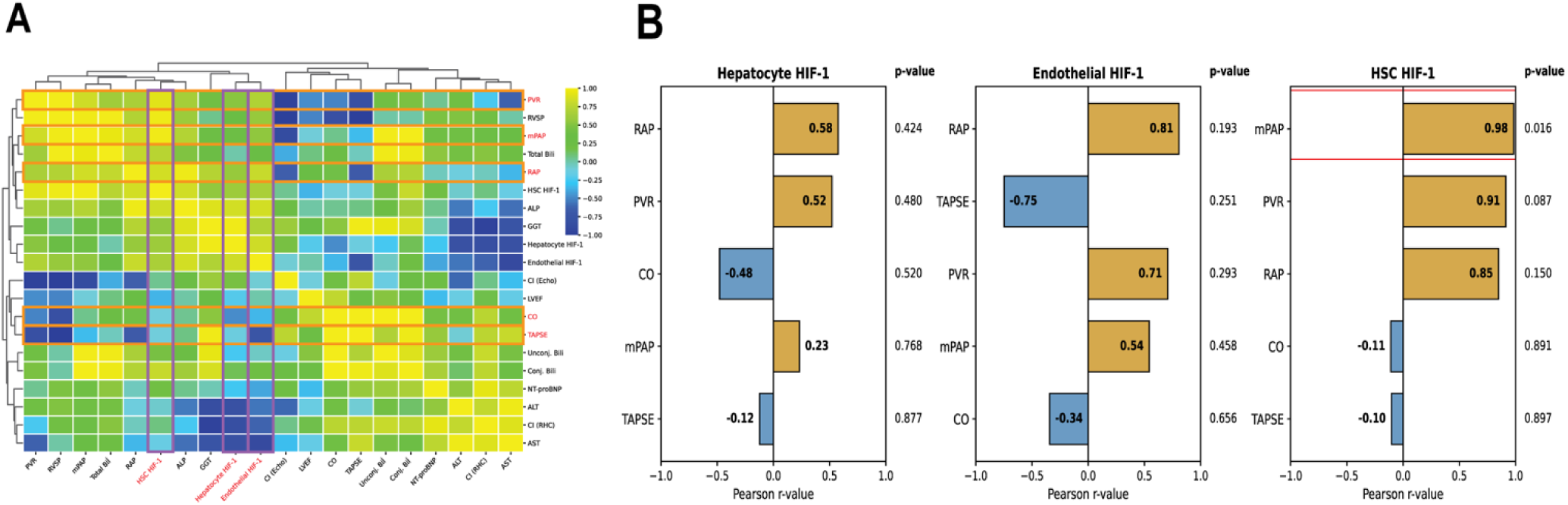
Correlational heatmapping identified relationships between right ventricular function and HIF-1 signaling in hepatocytes, endothelial cells, and hepatic stellate cells. (A) Heatmap displaying correlations between clinical markers of right ventricular failure/PAH severity and cell type-specific HIF-1 pseudobulked module scores in hepatocytes, endothelial cells, and hepatic stellate cells. (B) Correlation coefficients (r) and corresponding *p*-values for the top five associations for each cell type-specific HIF-1 module score.

**Supplemental Figure 10:**
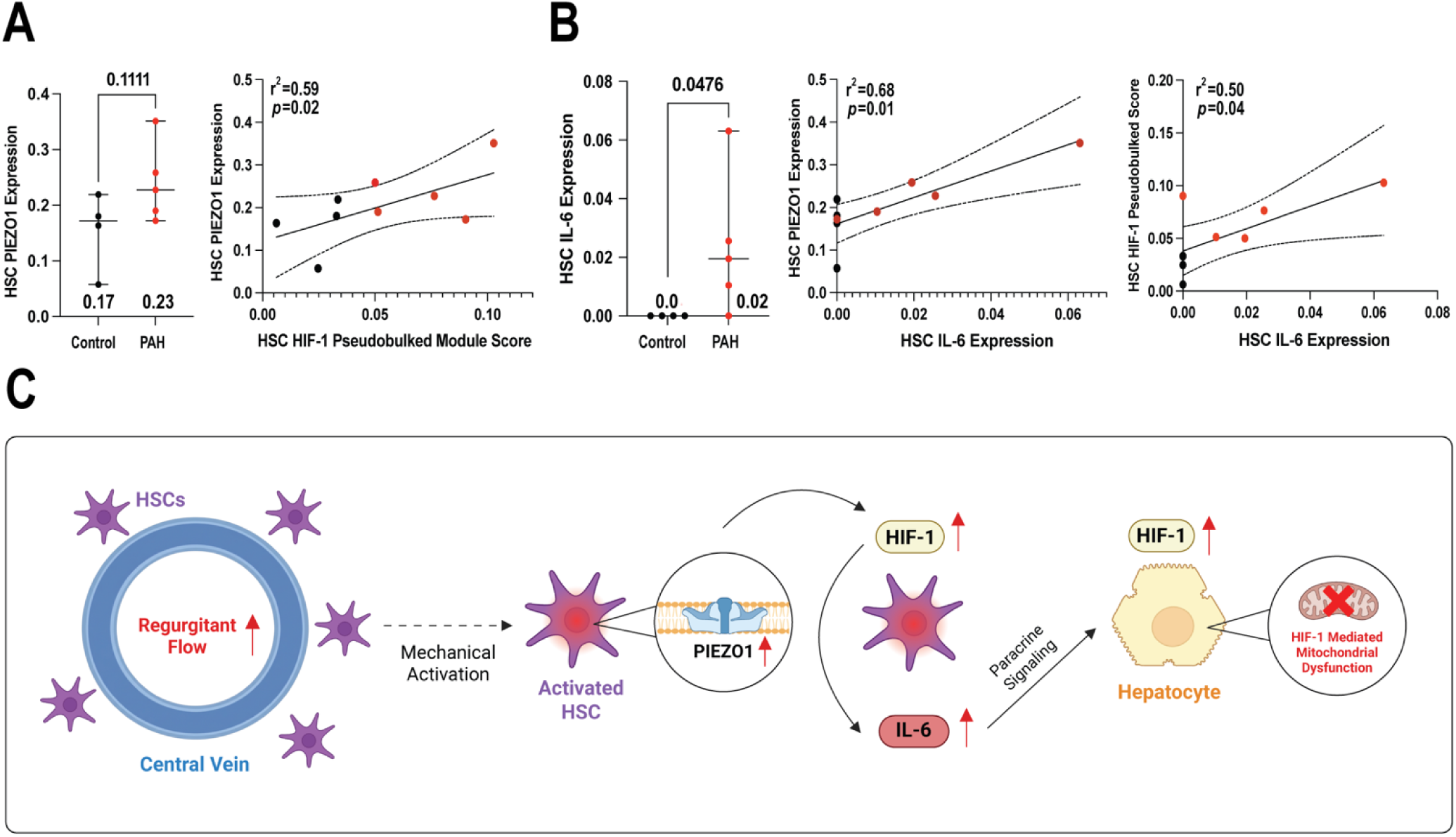
Hepatic stellate cell mechanosensitive and inflammatory signaling associate with HIF-1 activation. **(A)** HSC PIEZO1 expression in control and PAH livers and its relationship with HSC HIF-1 pseudobulked module scores. Dot plots display median with range (Mann-Whitney test) and correlation plots show linear regression with 95% confidence intervals. **(B)** HSC IL-6 expression in control and PAH livers and its associations with PIEZO1 expression and HIF-1 signaling. Dot plots display median with range (Mann-Whitney test) and correlation plots show linear regression with 95% confidence intervals. **(C)** Schematic illustrating the proposed mechanism linking HSC activation to downstream HIF-1 signaling.

**Supplemental Figure 11:**
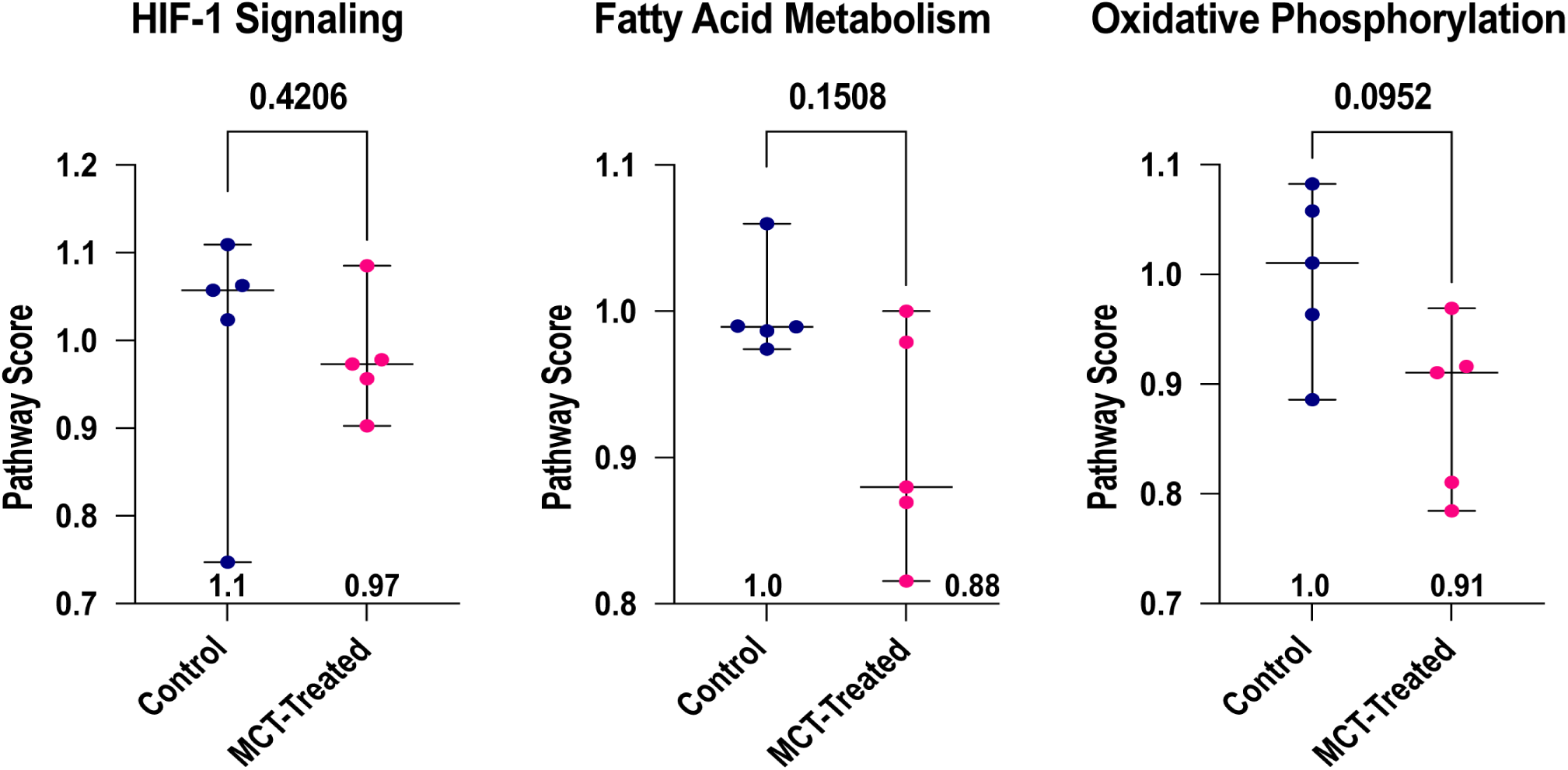
Monocrotaline (MCT)-treated rodent livers exhibit trends toward reduced fatty acid metabolism and oxidative phosphorylation, consistent with some of the metabolic alterations observed in human PAH livers. Error bars represent median with range, and *p*-values were calculated using Mann-Whitney tests.

