## Supplement for "Pulmonary Arterial Hypertension Induces a Metabolic and Inflammatory Hepatopathy"

**Supplemental Figure 1: Schematic representation of single nucleus RNA-sequencing workflow and data processing pipeline.**

**Supplemental Figure 2: Violin plots displaying key snRNA-seq quality control metrics across datasets, including total reads per nuclei, detected genes per cell, percent mitochondrial reads, and percent ribosomal reads.** Median values are displayed and error bars represent medians with interquartile range.

**Supplemental Figure 3: Sample-specific relative abundance values across cell types in PAH, NASH, and FALD datasets.** (A) Cell-type relative abundances in control and PAH samples. Error bars represent median values with range. (B) Cell-type relative abundances in control and NASH samples. Error bars represent median values with range. (C) Cell-type relative abundances in control and FALD samples. Error bars represent median values with range.

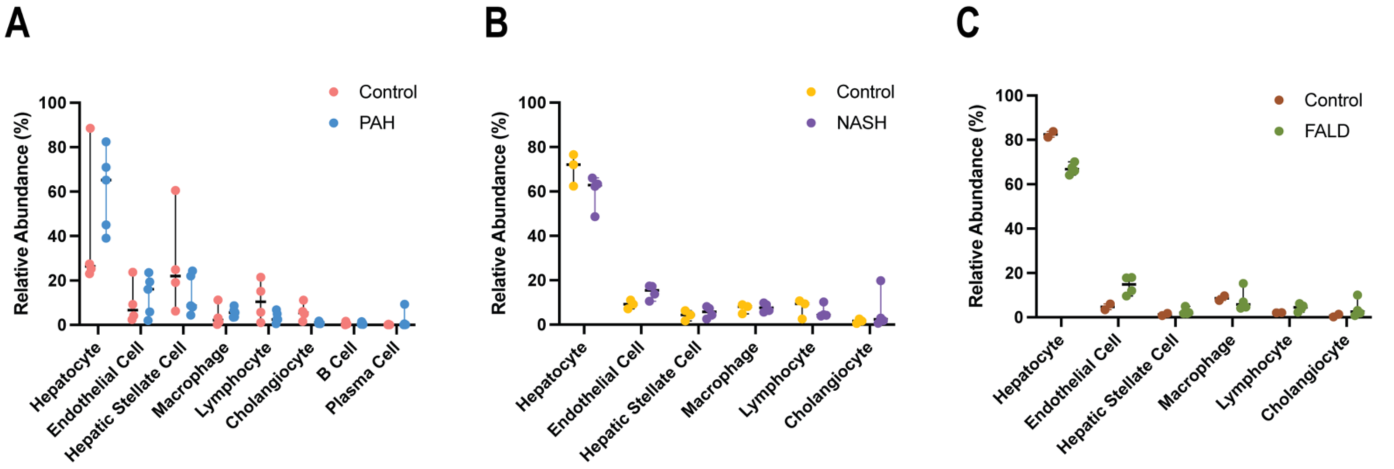

**Supplemental Figure 4: KEGG and Wiki pathway analysis of enriched and suppressed DEGs from PAH, NASH, and FALD hepatocytes relative to their respective controls.**

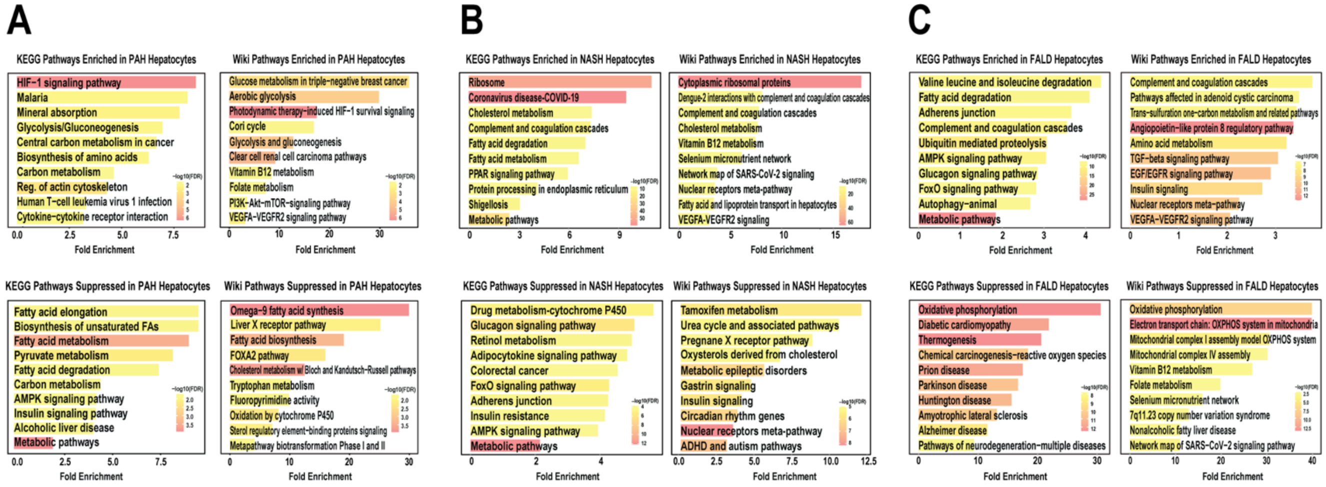

**Supplemental Figure 5: KEGG and Wiki pathway analysis of enriched and suppressed DEGs from PAH, NASH, and FALD endothelial cells relative to their respective controls.**

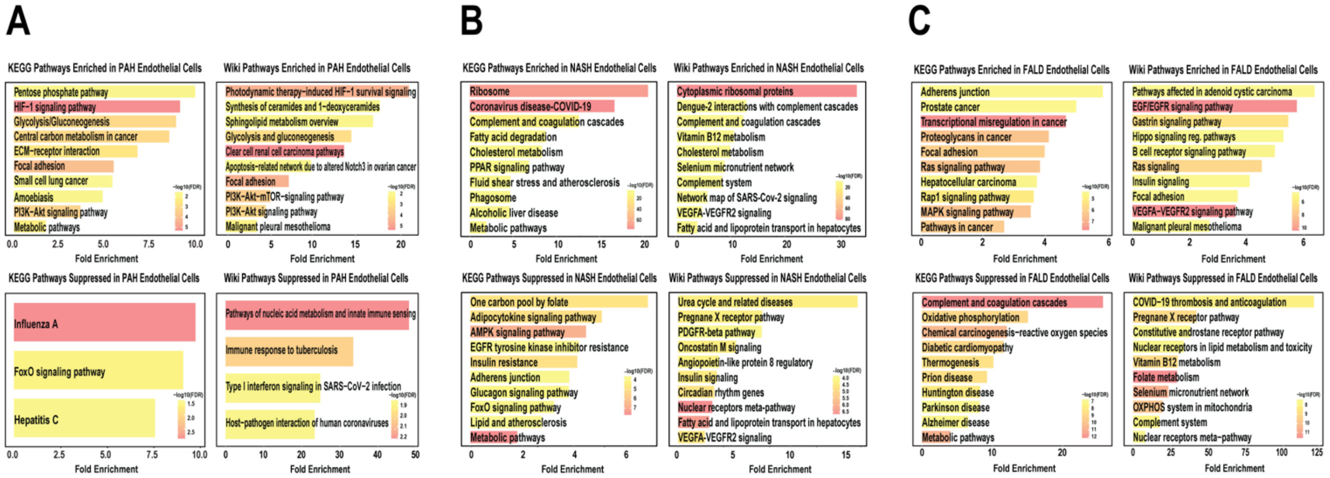

**Supplemental Figure 6: KEGG and Wiki pathway analysis of enriched and suppressed DEGs from PAH, NASH, and FALD hepatic stellate cells relative to their respective controls.**

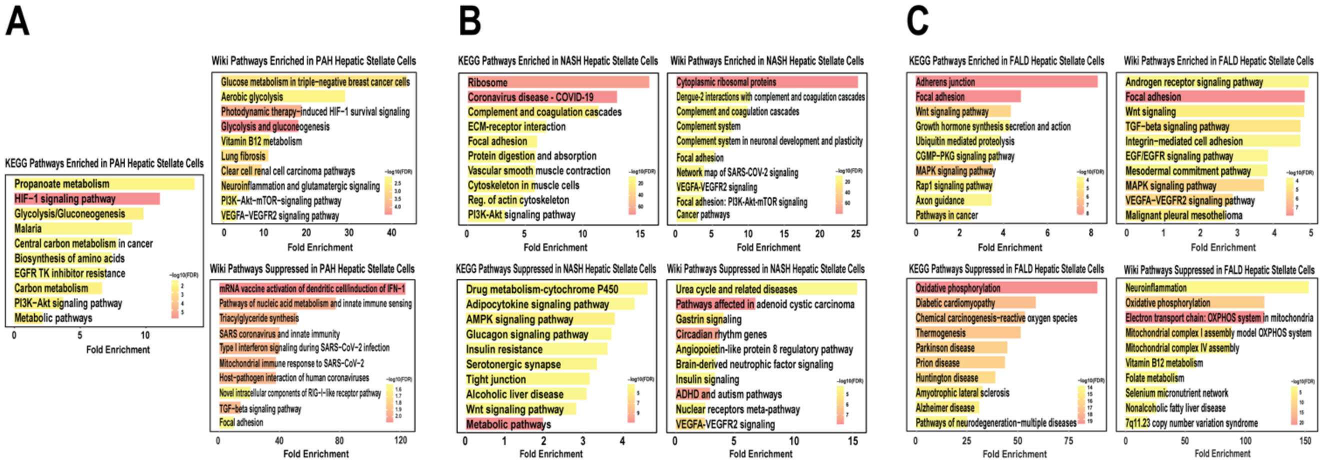

**Supplemental Figure 7: Total hepatic fibrosis was elevated in PAH livers and significantly associated with hepatic stellate cell HIF-1 signaling.** (A) Representative whole-liver images and quantification of trichrome total fibrosis (%) staining in control and PAH livers. Error bars represent median values with range, and the *p*-value was calculated by Mann-Whitney test. (B) Linear regression analysis showing the association between total fibrosis (%) and hepatic stellate cell HIF-1 pseudobulked module scores in control (black) and PAH (red) patients. Dotted lines display 95% confidence intervals.

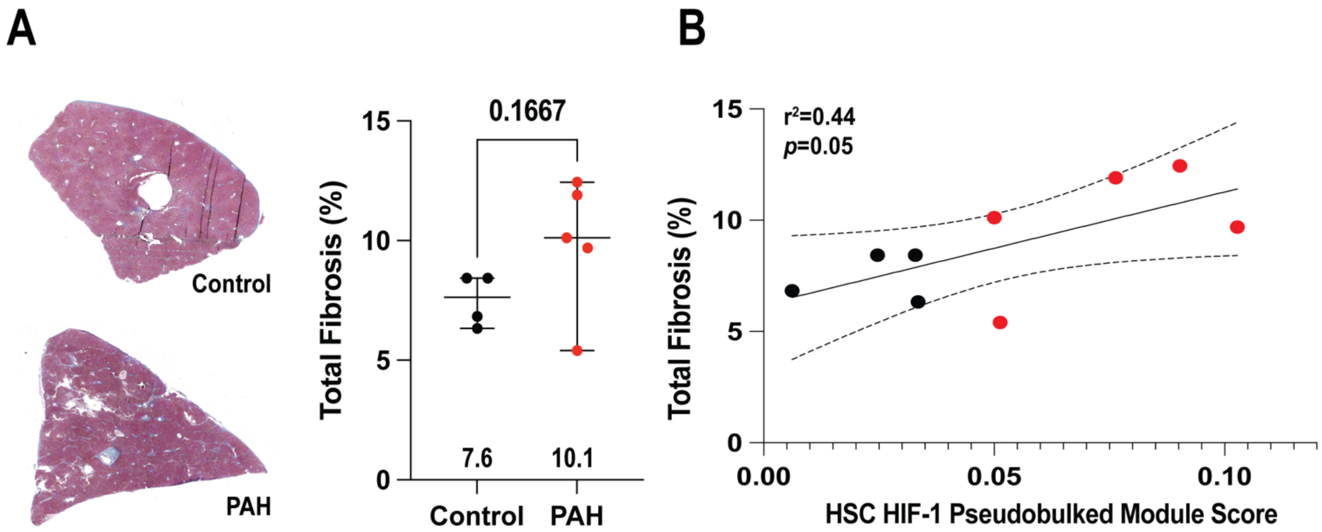

**Supplemental Figure 8: KEGG and Wiki pathway analysis of enriched and suppressed DEGs from PAH, NASH, and FALD macrophages relative to their respective controls.**

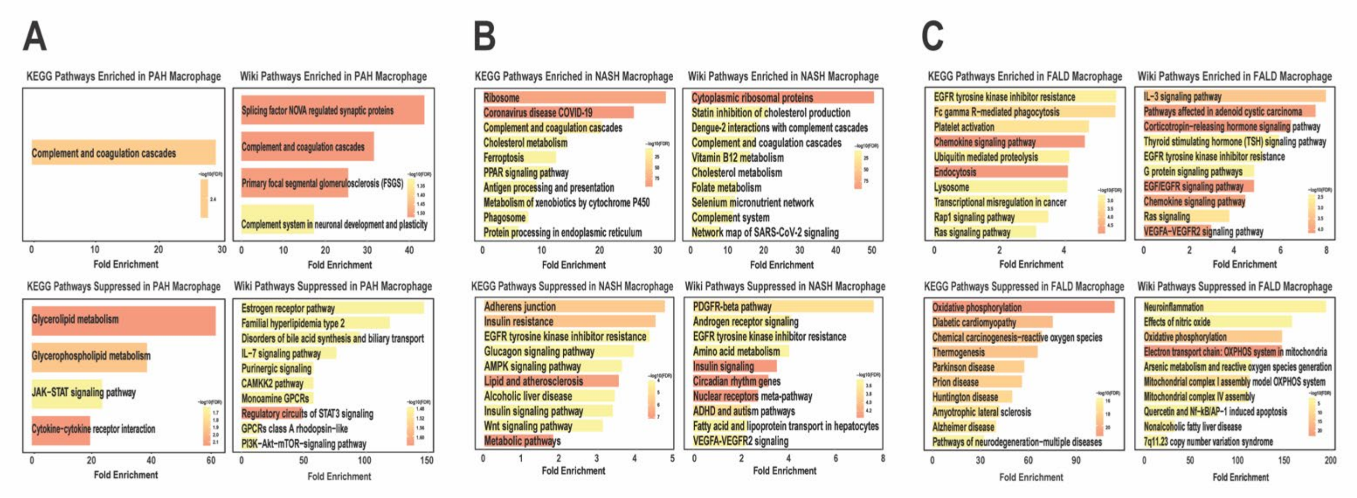

**Supplemental Figure 9: Correlational heatmapping identified relationships between right ventricular function and HIF-1 signaling in hepatocytes, endothelial cells, and hepatic stellate cells.** (A) Heatmap displaying correlations between clinical markers of right ventricular failure/PAH severity and cell type-specific HIF-1 pseudobulked module scores in hepatocytes, endothelial cells, and hepatic stellate cells. (B) Correlation coefficients (r) and corresponding *p*-values for the top five associations for each cell type-specific HIF-1 module score.

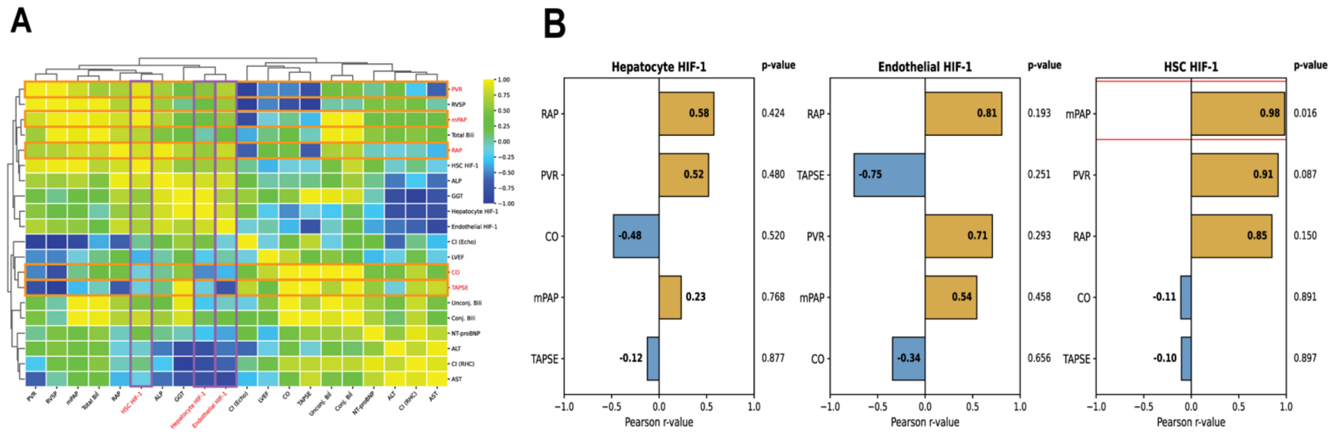

**Supplemental Figure 10: Hepatic stellate cell mechanosensitive and inflammatory signaling associate with HIF-1 activation. (A)** HSC PIEZO1 expression in control and PAH livers and its relationship with HSC HIF-1 pseudobulked module scores. Dot plots display median with range (Mann-Whitney test) and correlation plots show linear regression with 95% confidence intervals. **(B)** HSC IL-6 expression in control and PAH livers and its associations with PIEZO1 expression and HIF-1 signaling. Dot plots display median with range (Mann-Whitney test) and correlation plots show linear regression with 95% confidence intervals. **(C)** Schematic illustrating the proposed mechanism linking HSC activation to downstream HIF-1 signaling.

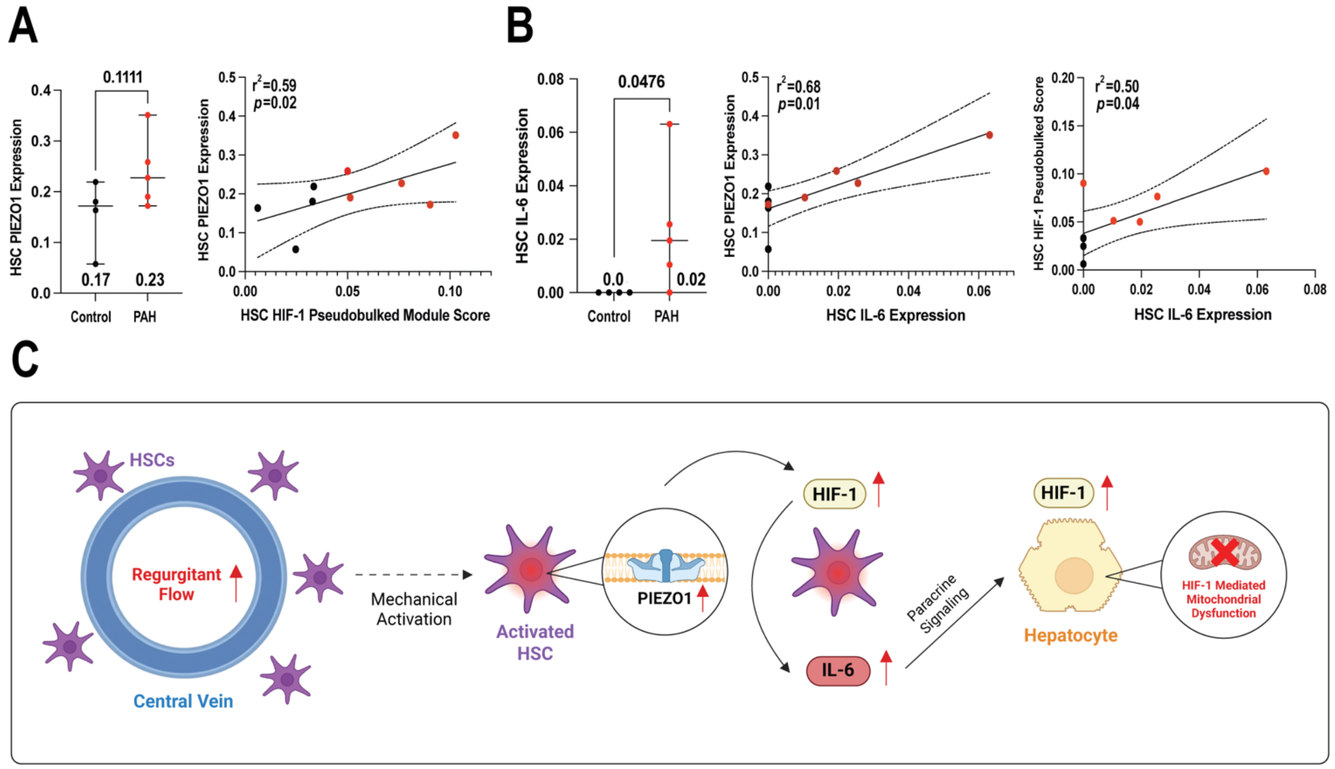

**Supplemental Figure 11: Monocrotaline (MCT)-treated rodent livers exhibit trends toward reduced fatty acid metabolism and oxidative phosphorylation, consistent with some of the metabolic alterations observed in human PAH livers.** Error bars represent median with range, and *p*-values were calculated using Mann-Whitney tests.

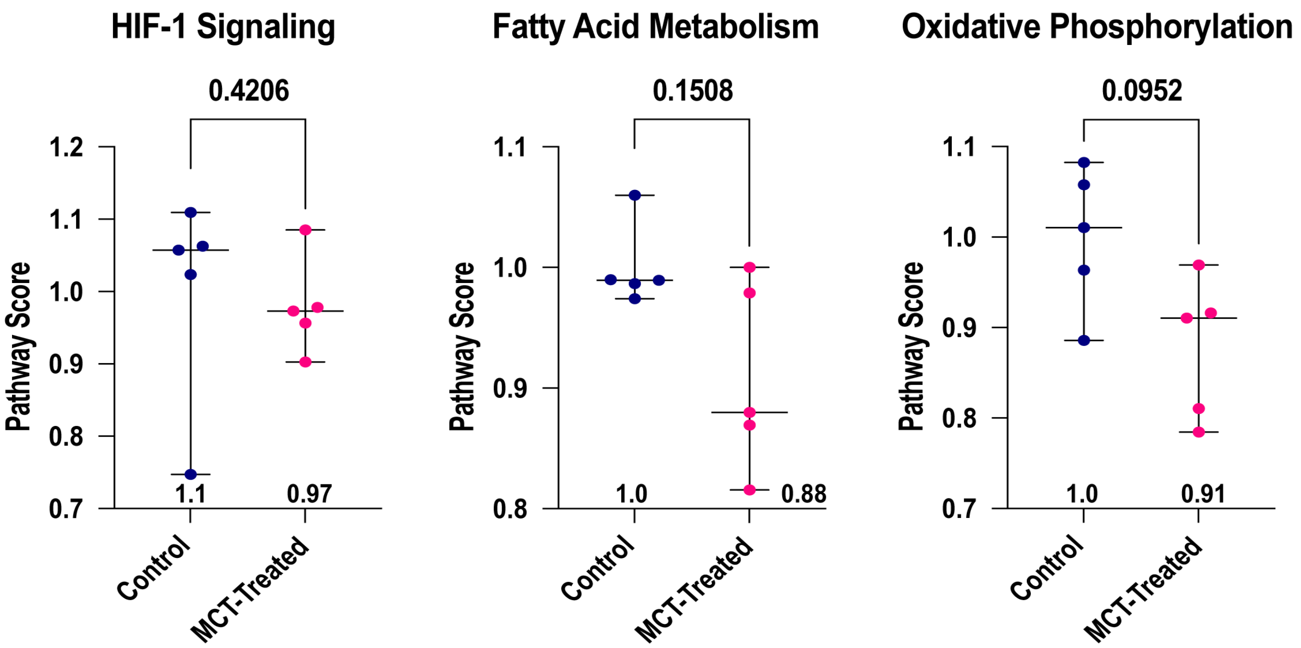
